# Bnip3lb-driven mitophagy sustains expansion of the embryonic hematopoietic stem cell pool

**DOI:** 10.1101/2024.09.23.614531

**Authors:** Eleanor Meader, Morgan T. Walcheck, Mindy R. Leder, Ran Jing, Paul J. Wrighton, Wade W. Sugden, Mohamad A. Najia, Isaac M. Oderberg, Vivian M. Taylor, Zachary C. LeBlanc, Eleanor D. Quenzer, Sung-Eun Lim, George Q. Daley, Wolfram Goessling, Trista E. North

## Abstract

Embryonic hematopoietic stem and progenitor cells (HSPCs) have the unique ability to undergo rapid proliferation while maintaining multipotency, a clinically-valuable quality which currently cannot be replicated in vitro. Here, we show that embryonic HSPCs achieve this state by precise spatio-temporal regulation of reactive oxygen species (ROS) via Bnip3lb-associated developmentally-programmed mitophagy, a distinct autophagic regulatory mechanism from that of adult HSPCs. While ROS drives HSPC specification in the dorsal aorta, scRNAseq and live-imaging of *Tg(ubi:mitoQC)* zebrafish indicate that mitophagy initiates as HSPCs undergo endothelial-to-hematopoietic transition and colonize the caudal hematopoietic tissue (CHT). Knockdown of *bnip3lb* reduced mitophagy and HSPC numbers in the CHT by promoting myeloid-biased differentiation and apoptosis, which was rescued by anti-oxidant exposure. Conversely, induction of mitophagy enhanced both embryonic HSPC and lymphoid progenitor numbers. Significantly, mitophagy activation improved ex vivo functional capacity of hematopoietic progenitors derived from human-induced pluripotent stem cells (hiPSCs), enhancing serial-replating hematopoietic colony forming potential.

**HIGHLIGHTS:**

1. ROS promotes HSPC formation in the dorsal aorta but negatively affects maintenance thereafter.
2. HSPCs colonizing secondary niches control ROS levels via Bnip3lb-directed mitophagy.
3. Mitophagy protects nascent HSPCs from ROS-associated apoptosis and maintains multipotency.
4. Induction of mitophagy enhances long-term hematopoietic potential of iPSC-derived HSPCs.

## INTRODUCTION

Compared to adult HSCs, embryonic HSCs have distinctive functional characteristics, including greater ability to proliferate while maintaining full multipotent erythro-myeloid and lymphoid potential, and naïve immunity allowing less rigorous matching for use in therapeutic transplantation^1^. This implies activation of specialized mechanisms for establishing these attributes unique to early development. Here, we show that spatio-temporally programmed mitophagy initiated by Bnip3lb, a pathway not utilized by adult HSCs, is key to the precise regulation of metabolic state that grants embryonic HSCs their unusual maintenance properties.

In both the embryo and the adult, Reactive Oxygen Species (ROS), byproducts of mitochondrial oxidative phosphorylation (OX/PHOS), act as key regulators of HSC fate. While normally contained within the mitochondrial outer membrane, overproduction or loss of membrane integrity allows for leakage of ROS into the cytoplasm, where it can damage other cellular components ^2^. This is referred to as oxidative stress^2^. Despite this negative attribute, ROS is an essential signaling molecule which influences the balance between hematopoietic proliferation and differentiation ^2–5^. In the adult bone marrow (BM), quiescent HSCs maintain low ROS and glycolytic metabolism^3,6,7^, then transition to OX/PHOS metabolism, and subsequent ROS production, when cycling and/or driving maturation.^8^ Although this switch is necessary for production of lineage restricted blood progenitors, excess and/or unregulated ROS leads to exhaustion of the HSC pool via unrestricted differentiation or apoptosis^9–18^. Conversely, if ROS production is completely inhibited, HSCs are prevented from cycling and producing downstream progenitors, resulting in an inability to fight infection or maintain hematopoietic homeostasis ^6,14,19,20^, particularly in the context of stem cell transplantation. Thus, metabolic state and ROS levels must be exquisitely balanced to support hematopoietic demand for the organism’s lifespan.^10,21^

Adult HSCs regulate ROS levels using a number of mechanisms, including maintaining quiescence, localizing within a hypoxic niche, utilizing glycolytic metabolism^14,15,22–26^ and employing high levels of “mitophagy” ^9,8,27–32^. The latter, a specialized form of cellular autophagy, is initiated when proteins on the outer membrane of a mitochondrion are tagged to mark the organelle for destruction.

Consequently, autophagy machinery constructs a membrane to surround the mitochondrion, which is then is fused to a lysosome, allowing contents to be digested and recycled. Broadly, there are two major pathways controlling mitophagy: the Pink1-Parkin pathway and receptor-mediated pathway, each of which can be regulated both transcriptionally and post-transcriptionally to fine-tune ROS levels within the cell^33^. Pink1-Parkin directed mitophagy occurs in tissues, including adult BM HSCs^11,19,34^, in response to damage and oxidative stress. Loss of outer membrane integrity leads to Pink1 translocation from the matrix to surface of the mitochondria. Pink1 is then ubiquitinated by Parkin, which draws in the autophagy machinery^35–37^. In contrast, receptor-mediated mitophagy occurs when the autophagy machinery is activated by mitophagy receptors, such as Fundc1, Bnip3, and Bnip3l (formerly Nix, which has two zebrafish homologues, Bnip3la and Bnip3lb), expressed on the surface of a mitochondrion^38,39^. Receptor mediated mitophagy is responsive not only to oxidative stress^40^ but also to hypoxia^41^and other developmental cues^33^; for example, Bnip3l/Nix is used to create space for hemoglobin in red blood cells,^42–44^ or minimize interference to light in developing corneal lens fibers^45–47^. While damage-responsive mitophagy is well documented in adult HSCs, there are no published roles for receptor-mediated mitophagy, although there is reportedly heterogeneity in autophagy pathways used by differentiating progenitors of various blood lineages^48^.

In contrast to the detrimental role generally ascribed to elevated ROS in the adult, our lab, and others, have shown that induction of ROS-mediated transcriptional activation is necessary for specification of HSPCs in the embryo. In this context, ROS elevation, coinciding with onset of blood flow and initiation of embryonic OX/PHOS metabolism^49^, stimulates a myriad of Hypoxia Inducible Factor 1a (Hifa)-associated gene targets critical for hemato-vascular development ^50–52^ and sterile inflammatory signaling via the NLRP3-inflammasome^49^. To explain this apparent shift in the regulatory impact of ROS on HSCs during early embryogenesis and their mature state, we must understand their developmental journey. After arising de novo from hemogenic endothelium (HE) in the ventral dorsal aorta (VDA), via an energetically demanding, metabolically active process termed endothelial-to-hematopoietic transition (EHT)^53^, HSCs immediately migrate to a secondary niche: the fetal liver (FL) sinusoids in mammals, or a vascular plexus in the tail known as the caudal hematopoietic tissue (CHT) in zebrafish^54^. There they proliferate to produce sufficient HSCs to fill the terminal niche and aid in populating subsequent waves of myeloid, lymphoid and erythroid lineages. Notably, during this “expansion phase”, FL HSCs utilize OX/PHOS metabolism more so than in the adult^55^. Following this window of exponential growth and differentiation, HSCs seed their terminal niche, the mammalian BM or kidney marrow (KM) in fish, where they take on adult characteristics, such as quiescence ^56,57^. The distinct environmental conditions of each niche, and unique activities undertaken by HSCs within them, presumably necessitate differential regulatory mechanisms to balance ROS, although little is known about use of mitophagy prior to BM colonization.

The work described herein documents the shifting spatio-temporal impact of ROS in the vertebrate embryo and elucidates the key regulatory role of developmentally programmed mitophagy in establishing the lifelong pool of HSPCs. Through scRNAseq and an in vivo fluorescent mitophagy reporter, we show that mitophagy is initiated in nascent HSPCs during EHT, with activation regulated by developmentally-programmed expression of the *bnip3lb* receptor. Gene knockdown studies confirm that embryonic induction of mitophagy is essential for maintenance of the self-renewing HSPC population within the CHT. Furthermore, antioxidant epistasis experiments indicate that mitophagy functions to protect the embryonic HSPC population by regulating ROS levels to balance maintenance of multi-potency while allowing for proliferative expansion in the absence of apoptosis. Finally, we reveal that activation of mitophagy via genetic or chemical stimulation, such as via Nicotinamide Riboside (NR), can significantly enhance production of HSPCs in vivo in the zebrafish embryo, as well as promote ex vivo maintenance of long-lived multipotent hematopoietic progenitors from human iPSCs.

## RESULTS

### ROS is essential for HSPC specification but detrimental as HSPCs leave the VDA niche

We previously found that elevation in ROS levels downstream of embryonic metabolic activation drives HSPC production in the VDA^49,50^. Given that sustained ROS stimulation negatively influences adult HSPC maintenance, we sought to understand at which developmental timepoint(s), and by what mechanism, the impact of ROS on embryonic HSPCs is regulated. To begin, we examined effects of ROS manipulation on HSPCs in the specification niche (the VDA), compared to the expansion niche (the CHT) (**Figure 1a**).

**Figure 1:**
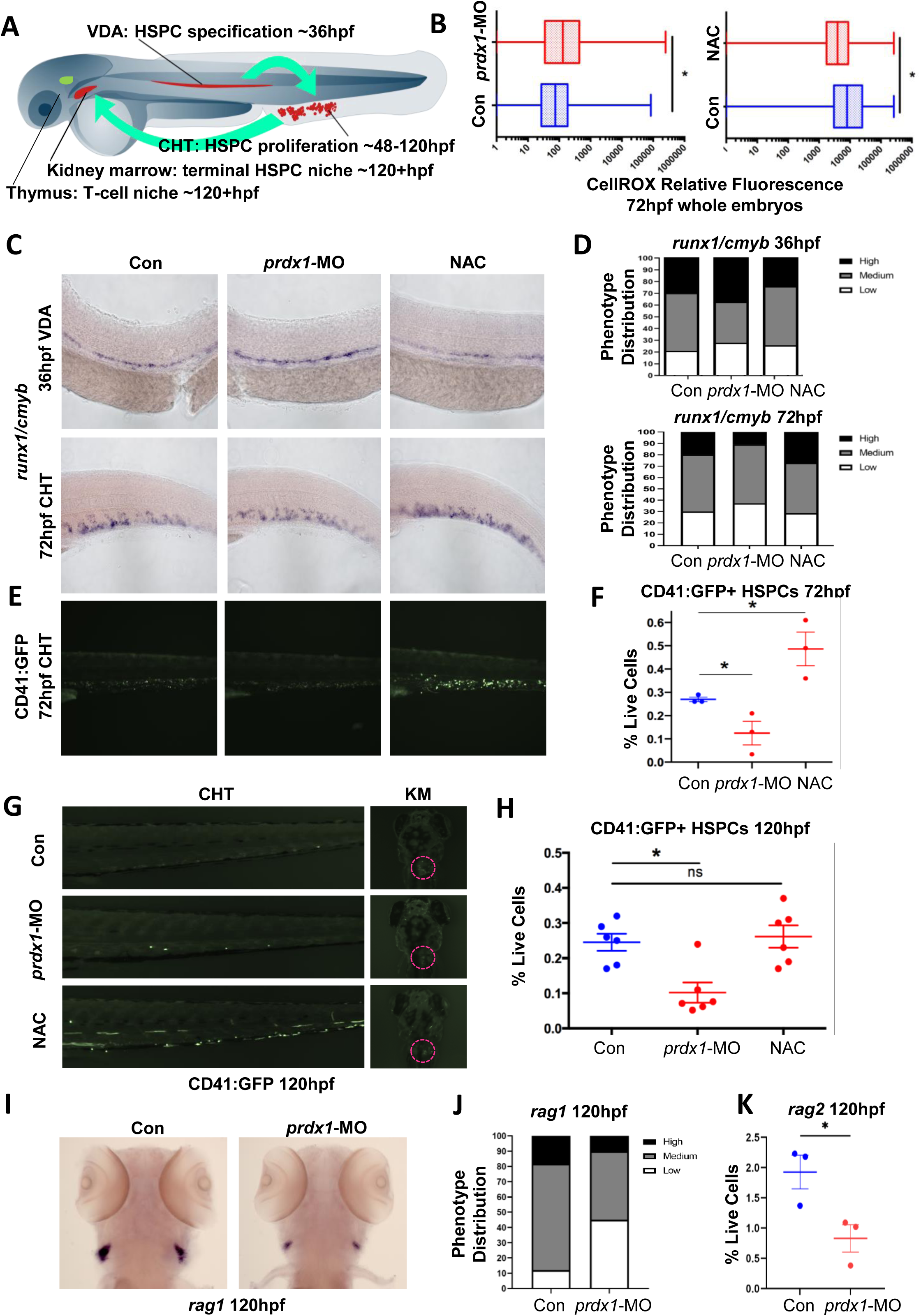
ROS is essential for HSPC specification, but detrimental following VDA exit (See also Figure S1) (A): Schematic indicating zebrafish hematopoietic niche sites (B): ROS levels in 72hpf embryos measured by flow cytometry using CellROX Orange^TM^ fluorescent ROS-dye. Representative examples of n=3 independent runs, significance calculated in FlowJo by K-S test (C): Representative images showing *runx1/cmyb* expression in embryos at 36hpf (10uM NAC Tx: 24–36 hpf; *top row*) and 72hpf (10uM NAC tx: 48-72hpf, *middle row*) assessed by WISH (D): Phenotypic distribution of *runx1/cmyb* expression scored as high, medium and low relative to sibling control embryos from (C) (E) *Tg(CD41:EGFP)* embryos (*bottom row*) were treated as above (C) and imaged at 72hpf (F) Flow cytometry for CD41:GFP+ HSPCs in *Tg(CD41:EGFP)* embryos at 72hpf after *prdx1* knockdown (p = 0.0487) or NAC treatment (p=0.0410). Mean±SEM, n=8 embryos/clutch, 3 clutches (G) *Tg(CD41:EGFP)* embryos treated with *prdx1*-MO or NAC (10uM, 48–120hpf), then imaged at 120hpf. KM = kidney marrow, outlined in magenta (H): CD41:GFP+ HSPCs by flow cytometry at 120hpf after *prdx1*-MO and NAC treatment (10uM, Tx: 48– 120hpf) (p=0.013, Mean±SEM, n=8 whole embryos/ clutch, 6 clutches) (I): Representative WISH images of *rag1* in the thymus at 120hpf (J): Phenotypic distribution of *rag1* expression scored in embryos from (I) (K): Flow cytometry for Rag2+ lymphocytes in Tg(rag2:gfp) embryos at 120hpf after *prdx1* knockdown (p=0.0378, Mean±SEM, n=8-10 embryos/clutch, 3 clutches,)

To elevate ROS levels in vivo, we microinjected morpholino oligonucleotides (MO) to knock down expression of *peroxiredoxin 1* (*prdx1)*, an enzyme which reduces hydrogen peroxide and other ROS^50^. Conversely, incubation with the antioxidant N-acetyl cystine (NAC; 10uM) was used to decrease ROS levels, as previously described^50^. In vivo alterations in ROS levels were evaluated by flow cytometry of disaggregated 48hpf embryos using CellROX, a dye which fluoresces in the presence of ROS (**Figure 1b**). As in our prior report^50^, whole mount in situ hybridization (WISH) showed increased ROS promoted expression of the HSPC markers *runx1* and *cmyb* in the VDA at 36hpf (**Figure 1c, d, *top row*)**. However, by 72hpf, after HSPCs have migrated to the CHT, continued ROS elevation in *prdx1* morphants reduced *runx1/cmyb* expression, as shown in the representative images and graphic summary of the qualitative phenotypic evaluation of total embryos scored per condition (**Figure 1c,d, *middle row***). Similar biphasic alterations in HSPC markers were observed when increasing ROS by H_2_O_2_ or 1% glucose treatment (**Supp. Figure 1a, b**). By contrast, as opposed to its negative impact on HSPC formation in the VDA (**Figure 1 c, d, *top row***), later incubation with the antioxidant NAC increased the appearance of HSPC markers in the CHT at 72hpf (**Figure 1c, d, *middle row***). This developmental flip in the impact of ROS on HSPCs in the VDA and CHT was validated by in vivo imaging using *Tg(CD41:EGFP)* HSPC-reporter embryos (**Figure 1e**), and further quantified by measuring the CD41:GFP^lo^ HSPC population via flow cytometry^58,59^, confirming both a significant increase in the NAC-treated HSPC population and a significant decrease in the *prdx1* morphant HSPC population at 72hpf (**Figure 1f**). Interestingly, the impact of ROS modulation at 48hpf, a timepoint at which HSPCs continue to bud in the AGM, while simultaneously colonizing the CHT, is more ambiguous (**Supp. Figure 1a, b, *middle rows***), reflecting a transitional phase for both HSPC localization and ROS regulation.

To assess whether ROS additionally impacts the HSPC population at later stages of development, and to address potential effects on HSPC-dependent lineage commitment, we analyzed embryos at 120hpf. Knockdown of *prdx1* significantly reduced the expression of the HSPC marker CD41 in the kidney marrow (**Figure 1g, h**) and lymphoid marker *rag1* in the thymus (**Figure 1i,j,k**); the latter observation was confirmed by flow cytometry for *Tg(rag2:GFP)*, showing a significant decrease at 120hpf (**Figure 1k**). Notably, *prdx1* morphants showed little change in expression of myeloid marker *mpo* at this timepoint by WISH or flow cytometry (**Supp. Figure 1c-e**), indicative of primary production from earlier hematopoietic waves^60^. Interestingly, delayed initiation of NAC treatment, from 48-120hpf, had little effect on CD41 expression in the KM (**Figures 1g, h**), presumably reflective of native activation of ROS limiting and/or regulatory factor(s) by this stage. Together, these data suggest ROS levels must be precisely controlled in HSPCs as embryonic development progresses to properly seed terminal niches and maintain balanced differentiation capacity.

### scRNASeq analysis shows *bnip3lb* expression increases in developing HSPCs

To understand how ROS transitions from boosting HSPC numbers in the VDA to reducing them in the CHT and KM, we examined these populations via single cell RNA-seq. Using *Tg(kdrl:Has.HRAS-mCherry)^s916^* vascular reporter embryos, we microdissected and disaggregated the trunk/tail region, then collected the Kdrl+ population, inclusive of hemogenic endothelium and newly formed HSPCs, at 36hpf by FACS. After sequencing the sorted population, we identified several distinct clusters, including one enriched for the expression of classical HSC markers (**Figure 2a, b**). Analysis of this cluster indicated that it fell into two distinct sub-clusters: one of which had prominent expression of endothelial (e.g. *fli1a, zeb2a, gfi1b and meis1b*) markers, leading us to deem it the “hemogenic endothelial cell” population (HEC cluster), while the other was more enriched for conserved HSC markers (*cmyb, spi2, ikzf1,* and *cxcr4b*), defining it as a more mature “HSPC-like” population (HSPC cluster)(**Figure 2c-e**).

**Figure 2:**
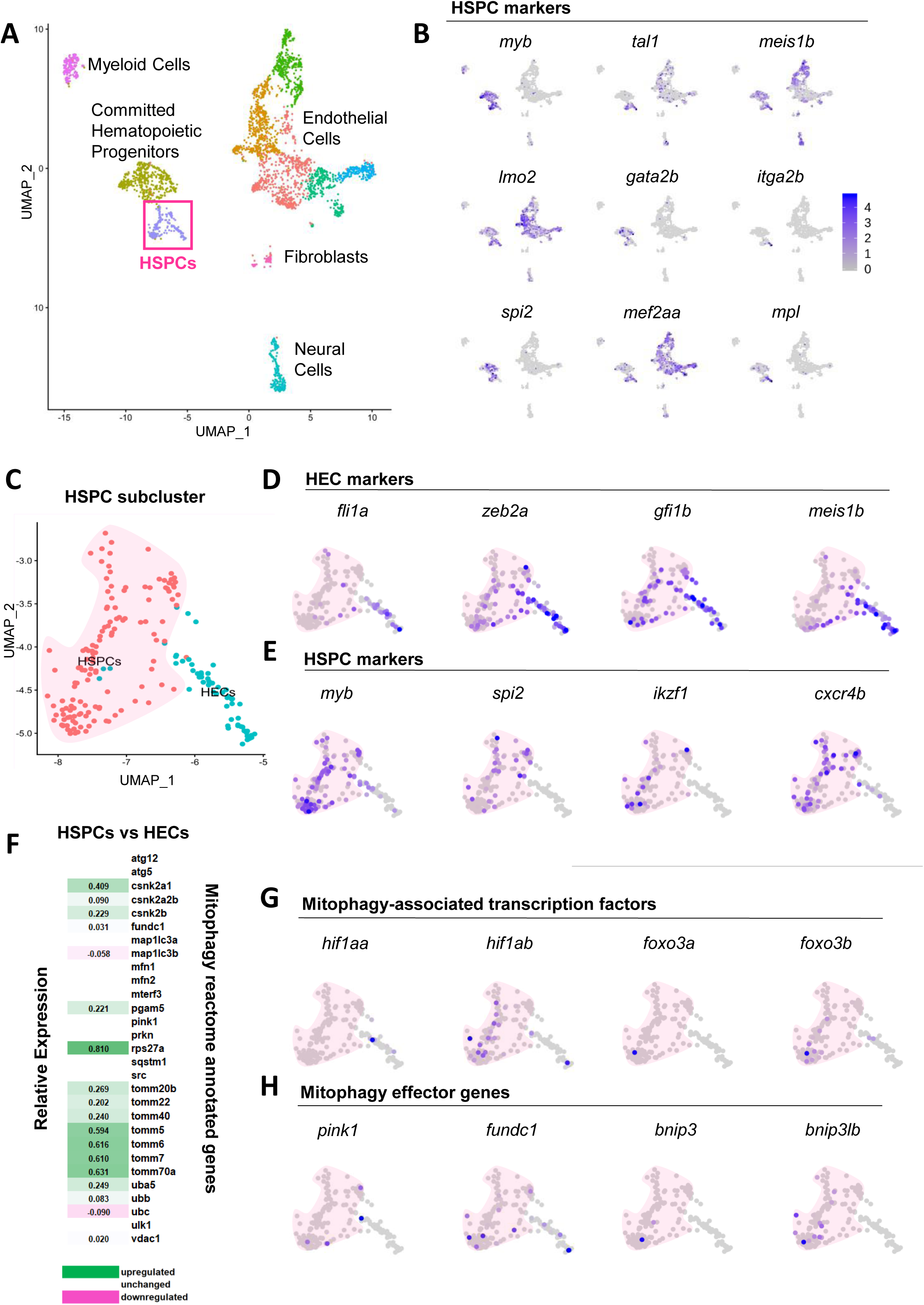
scRNASeq analysis reveals *bnip3lb* expression in nascent HSPCs (See also Figure S2) (A): UMAP-clustering of all sequenced Kdrl:mCherry+ cells sorted from dissociated tails of 36hpf WT embryos (B): UMAP expression patterns of select HSPC marker genes, revealing identity of an HSPC cluster (C): UMAP of isolated HSPC cluster (from magenta box in 2A), showing 2 sub-clusters labeled HECs (hemogenic endothelial cells) and HSPCs (hematopoietic stem and progenitor cells, *pink cloud*) (D): UMAP expression patterns of select HEC marker genes defining identity of the HEC sub-cluster (E): UMAP expression patterns of select HSPC marker genes demonstrating identity of maturing HSPC sub-cluster (F): Heatmap showing differential expression of MITOPHAGY_REACTOME annotated genes^106^ between the HEC and HSPC sub-clusters. Genes upregulated in the HSPC sub-cluster = green; genes downregulated = magenta. Genes not differentially regulated or without detectable expression = white. (G): UMAP showing expression patterns of select mitophagy-promoting transcription factors (H): UMAP showing expression patterns of select mitophagy pathway genes

Gene ontology (GO) analysis of annotated biological processes differentially expressed between these two populations revealed the HSPC sub-cluster was enriched for GO terms related to biosynthesis, perhaps reflecting the beginning of their rapid expansion phase; notably, this subcluster also showed significant enrichment for terms reflective of cellular stress, and in particular mitophagy (**Supp. Figure 2**). In targeted analysis of differentially expressed genes between the two subclusters, we noted upregulation of mitophagy-associated genes in the HSPC subcluster compared to the HEC population (**Figure 2f**). To better understand this correlation, we visualized expression of classical mitophagy-promoting transcription factors ^61,62^ and found a strong bias toward increased expression in the HSPC fraction (**Figure 2g**). To determine which pathways may be responsible for mediating the putative mitophagy signature in the HSPC population, we examined effector gene expression: whereas *parkin* and *bnip3la* were not expressed at detectable levels, *pink1* and *fundc1* were expressed at low levels in both subclusters. In contrast, *bnip3* and *bnip3lb* were uniquely differentially expressed between the HSPC and HEC populations, with higher expression in HSPCs, suggestive of directed upregulation during or immediately after EHT (**Figure 2h**).

### *Bnip3lb* is expressed in the HSPCs within the CHT, coinciding with increased mitophagy

To further document the spatio-temporal onset of mitophagy as it relates to definitive hematopoiesis, we utilized a recently-developed fluorescent mitophagy-reporter zebrafish line, *Tg(ubi:mitoQC)* ^63^. This tool, based on the mitoQC fluorescent cassette by Zhao et al.^64^, expresses both GFP and mCherry on the surface of mitochondria, allowing for live imaging of mitophagy. Under normal circumstances the GFP signal is brighter; however, when mitochondria undergo mitophagy, GFP is quenched by the low pH of the fused lysosome, and mCherry can be visualized (**Figure 3a**). After crossing *Tg(ubi:mitoQC)* to a *Tg(kdrl:bfp)* endothelial reporter line, mitochondrial dynamics of EHT were examined by live whole mount imaging at 36hpf (**Figure 3b, Supp Figure 3a**). As anticipated given the previously delineated role of BNIP3l-mediated mitophagy in erythroid maturation^42,43^, strong mCherry signal was detected in red blood cells in circulation in the dorsal aorta and posterior cardinal vein (PCV). Lower levels of mitophagy were also visible throughout the vasculature, with comparatively higher mCherry signal present in a discrete subset of Kdrl+ cells, which based on their rounded morphology and position at the VDA/PCV junction, were likely newly specified HSPCs. Indeed, timelapse imaging revealed novel mitophagy puncta emerging immediately after select Kdrl+ cells appeared to undergo EHT, culminating in budding from the arterial wall and migration into the interluminal space or circulation (**Figure 3b, additional images in Supp. Figure 3b**). Using the reporter, we further demonstrated significantly more mitophagy present in Kdrl+ cells of the CHT compared to the VDA, presumably inclusive of the Kdrl-derived nascent HSPCs (**Figure 3c**). To further validate this observation, we examined a *Tg(CD41:mitoQC)* line, confirming high levels of mitophagy localized in HSPCs present within the CHT niche (**Supp. Figure 3c**).

**Figure 3:**
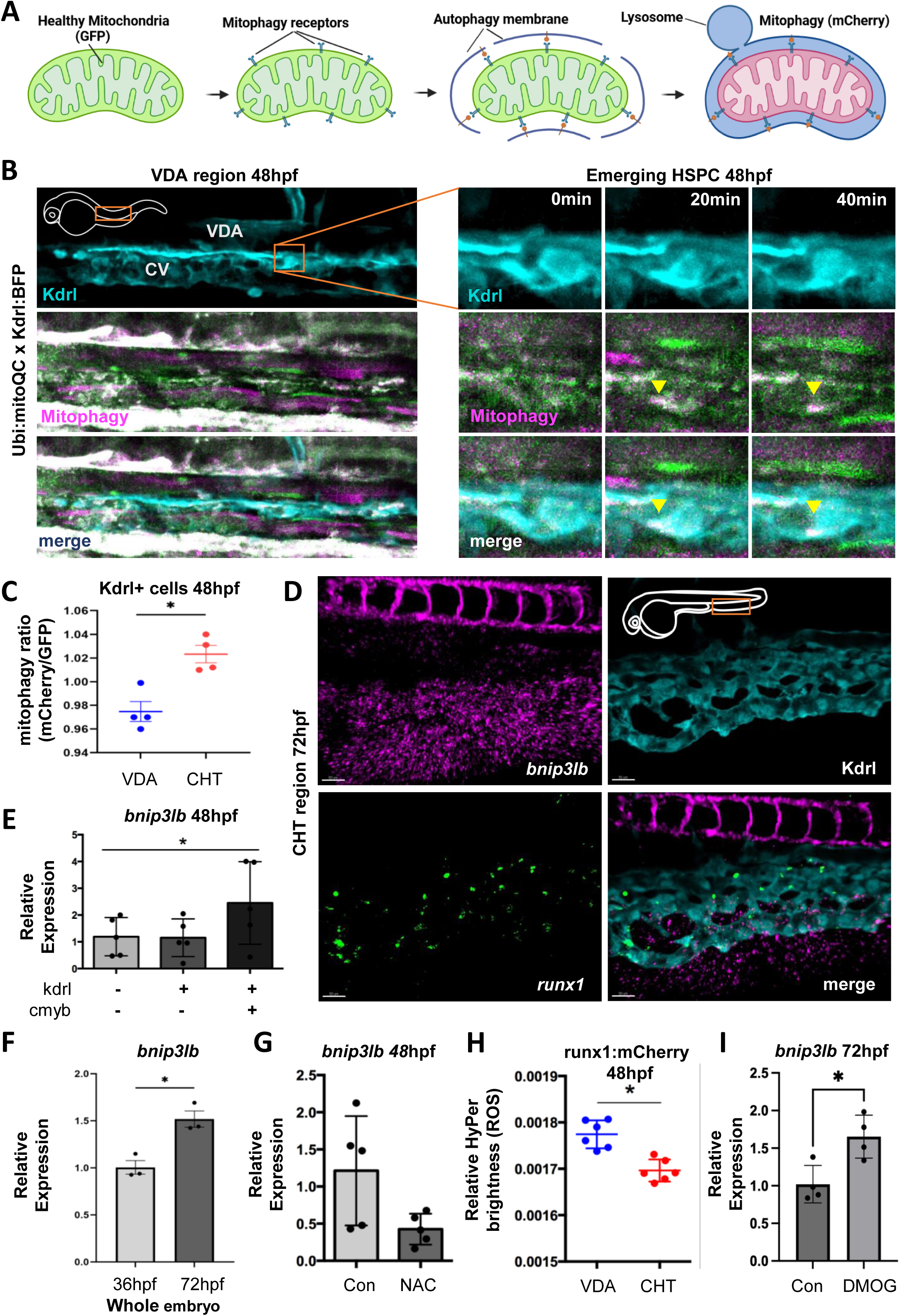
*Bnip3lb* expression in CHT HSPCs coincides with increased mitophagy (See also Figure S3) (A): Schematic of mitophagy reporter zebrafish model. Image created from Biorender.com (B): *Left*: Confocal image of mitophagy occurring in the VDA region (*orange box in embryo diagram inset*) at 48hpf. Healthy mitochondria (*green*), mitophagy (*magenta*), Kdrl+ vasculature (*blue*). *Right*: Timelapse of zoomed in image of mitophagy puncta (yellow arrow) forming in a budding HSPC. CV = Caudal Vein. (C): Ratiometric quantification of mitophagy (mCherry/GFP ratio) in Kdrl+ cells measured from images of *Tg(ubi:mitoQC;kdrl:BFP)* embryos at 48hpf (p=0.0048, Mean±SEM, n=4 embryos/condition) (D): Fluorescent WISH for *bnip3lb* expression (*magenta*) in the CHT at 72hpf, overlaid with WISH for *runx1* (HSPC, *green*) and antibody staining for Kdrl (vasculature, *blue*), representative of 5 larvae (E): *bnip3lb* qPCR on 48hpf FACS isolated Kdrl+cMyb+ HSPCs and Kdrl1+ vasculature normalized to the Kdrl-cMyb-fraction. (Mean±SEM, n=5, ANOVA p=0.0331) (F): qPCR for *bnip3lb* expression in 36hpf versus 72hpf embryos. p=0.0105, Mean±SEM (n=20 embryos/clutch, 3 clutches) (G): *bnip3lb* qPCR on 48hpf embryo tail fractions after treatment with 10uM NAC from 36-48hpf (p=0.0508, Mean±SEM, n= 5embryos/clutch, 5 clutches) (H): Quantification of ROS in Runx1:mCherry+ cells in the VDA vs CHT of 48hpf embryos, measured via the relative fluorescence of HyPer in confocal images (p=0.002, Mean±SEM, n=6) (I): *bnip3lb* qPCR on 72hpf tail fractions treated with Hif1a activator DMOG (10uM) from 36-48hpf (p=0.0160, Mean± EM, n=12-15 embryos/clutch, 6 clutches)

These results collectively suggest that mitophagy increases as HSPCs emerge via EHT and colonize the CHT, consistent with the scRNAseq data. As our bioinformatics analysis of the HSPC subcluster pointed to Bnip3lb as a promising mechanistic candidate to drive mitophagy during this transition, we sought to validate its expression in the nascent HSPC population. Cellular localization of *bnip3lb* was assessed utilizing fluorescent WISH (FISH), demonstrating co-expression with *runx1* within the CHT vascular plexus (**Figure 3d**). Using *Tg(cmyb:EGFP;kdrl:Has.HRAS-mCherry)* reporter embryos, qPCR was performed on FACS purified Kdrl-/cMyb- (control), Kdrl+/cMyb- (endothelial) and Kdrl+/cMyb+ (HSPC) cell populations at 48hpf (**Figure 3e**), further validating enrichment of *bnip3lb* in HSPCs in the CHT. Further, whole embryo qPCR analysis demonstrated that *bnip3lb* expression increased significantly between 36hpf and 72hpf (**Figure 3f**), coinciding with the transition of HSPCs from the VDA to the CHT. Significantly, this induction of *bnip3lb* expression coincided directly with initiation of mitophagy reporter expression in vivo, suggestive of developmentally-directed activation as HSPCs mature.

Whereas many mitophagy-associated effector genes are primarily regulated post transcriptionally in response to immediate stressors such as oxidative stress, starvation or hypoxia, our data indicate transcriptional initiation of *bnip3lb* during embryonic hematopoiesis. Bnip3l, or Nix, has an established role in developmentally programmed mitophagy, with expression regulated by metabolism-responsive transcription factors such as FOXO3, which controls expression of numerous ROS-response genes^61^, and HIF1a, which regulates a variety of genes in response to hypoxia^62^. In our RNASeq data, both *hif1ab* and *foxo3b* are expressed in a similar pattern to *bnip3lb* (**Figure 2g**), exhibiting enrichment in the HSPC cluster. We therefore knocked down each potential regulator to determine if it would impact expression of *bnip3lb* at timepoints relevant to HSPC development. Although not significant, *foxo3b* morphants trended toward increased *bnip3lb* expression by qPCR at 72hpf (**Supp. Figure 3d**), potentially reflective of redundancy in oxidative stress-responsive transcriptional and/or post-translational regulation of *bnip3l* expression. Mitophagy in general responded to ROS elevation at this stage of development, as shown by the mitoQC ratio after *prdx1* knockdown (**Supp. Figures 3e, f**); additionally, *bnip3lb* expression itself showed a modest trend upwards in *prdx1* morphants (**Supp. Figure 3g**), and a downward in response to NAC (10uM, from 36 to 48hpf) (**Figure 3g**) by qPCR at 48hpf, indicating it can be regulated in response to ROS. Importantly, by confocal mitophagy of the *Tg(ubb:HyPer) in vivo* ROS reporter line, ROS levels were decreased in Runx1+ HSPCs of the CHT compared to the VDA at 48hpf (**Figure 3h, Supp. Figure 3h**), correlating with increased *bnip3lb* expression and mitophagy activation. This suggests additional factors can promote expression of *bnip3lb* in HSPCs of the CHT. *hif1ab* is the primary Hif1 paralog driving HSC formation in zebrafish^52^; MO-mediated knockdown did not reduce *bnip3lb* expression by qPCR on tail isolates of 72hpf embryos (**Supp. Figure 3i**), however, activation of Hif1a with chemical hypoxia mimic DMOG^65^ (10uM, 48-72hpf) significantly increased *bnip3lb* expression the CHT at 72hpf (**Figure 3i**). This implies that developmentally programmed mitophagy in HSPCs is redundantly regulated by HIF, ROS and other stress-responsive pathways.

### *Bnip3lb* knockdown reduces embryonic HSPC and lymphoid populations after EHT

Given the developmentally synchronized induction of mitophagy and *bnip3lb* expression, we sought to determine if Bnip3lb-associated mitophagy is necessary for establishment and/or maintenance of the embryonic HSPC pool. Using a previously validated *bnip3lb*-MO^63^, we selected a dose with no observable effect on vasculature or primitive erythrocyte development (**Supp. Figure 4a, b**) to examine impact on the HSPC population. Knockdown of *bnip3lb* significantly reduced mitophagy reporter signal in the trunk/tail region (**Figure 4a**). Consistent with the developmental timing of *bnip3lb* expression and mitophagy reporter activity, HSPC marker expression in the VDA at 36hpf was not altered in morphants (**Figure 4b, c**), however, by 72hpf *bnip3lb* proved essential for maintaining the *runx1/cmyb*+ HSPC population in the CHT (**Figure 4d, e**). This observation was quantified using *Tg(CD41:EGFP)* embryos, which exhibited a significant loss in CD41+ HSCs by flow cytometry (**Figure 4f**). Interestingly, at 48hpf during the VDA to CHT transition, *runx1/cmyb* expression in the VDA was enhanced in *bnip3lb* morphants (**Supp. Figure 4c, d**), in line with prior reports^50^, indicating that elevated ROS can continue to increase HSPCs undergoing EHT at this timepoint and/or suggesting that initiation of mitophagy may play a role in HSPC egress or migration to the CHT. Importantly, this effect appeared to be specific for Bnip3lb-associated mitophagy, as MO-mediated *pink1* knockdown^66^ had no impact in HSPC gene expression at 72hpf (**Figure 4g, h**). Evaluation of hematopoietic development at later embryonic timepoints revealed that reduced mitophagy in *bnip3lb* morphants was associated with decreased expression of *rag1* in the thymus at 120hpf, and a corresponding increase in *mpo* (**Figure 4i-l**) in the CHT, suggestive of lineage skewing or failure of progenitors to migrate to or effectively seed subsequent niches. To verify these findings, a previously established *bnip3lb* mutant fish line^63^ was examined: mutants showed an unexpected increase in HSPCs in the CHT by *runx1/cmyb* WISH at 72hpf, however, this effect was strongly mitigated by combinatorial injection of morpholinos against the *bnip3lb* homologue *bnip3la* (**Supp. Figure 5a, b**); mutant embryos also demonstrated significant upregulation of other mitophagy related genes (**Supp Figure 5c**), suggesting the absence of an HSPC effect in *bnip3lb* mutants is likely due to genetic compensation, as reported for other gene families^67–69^. Although mitophagy can be regulated by multiple pathways with complex and compensatory mechanisms, chemically inhibiting autophagy via blocking lysosome fusion using bafilomycin (50uM), mimics *bnip3lb* knockdown, causing a dramatic reduction in *runx1/cmyb* expression in the CHT (**Supp. Figure 5d-g**), emphasizing the importance of mitophagy in this secondary hematopoietic niche.

**Figure 4:**
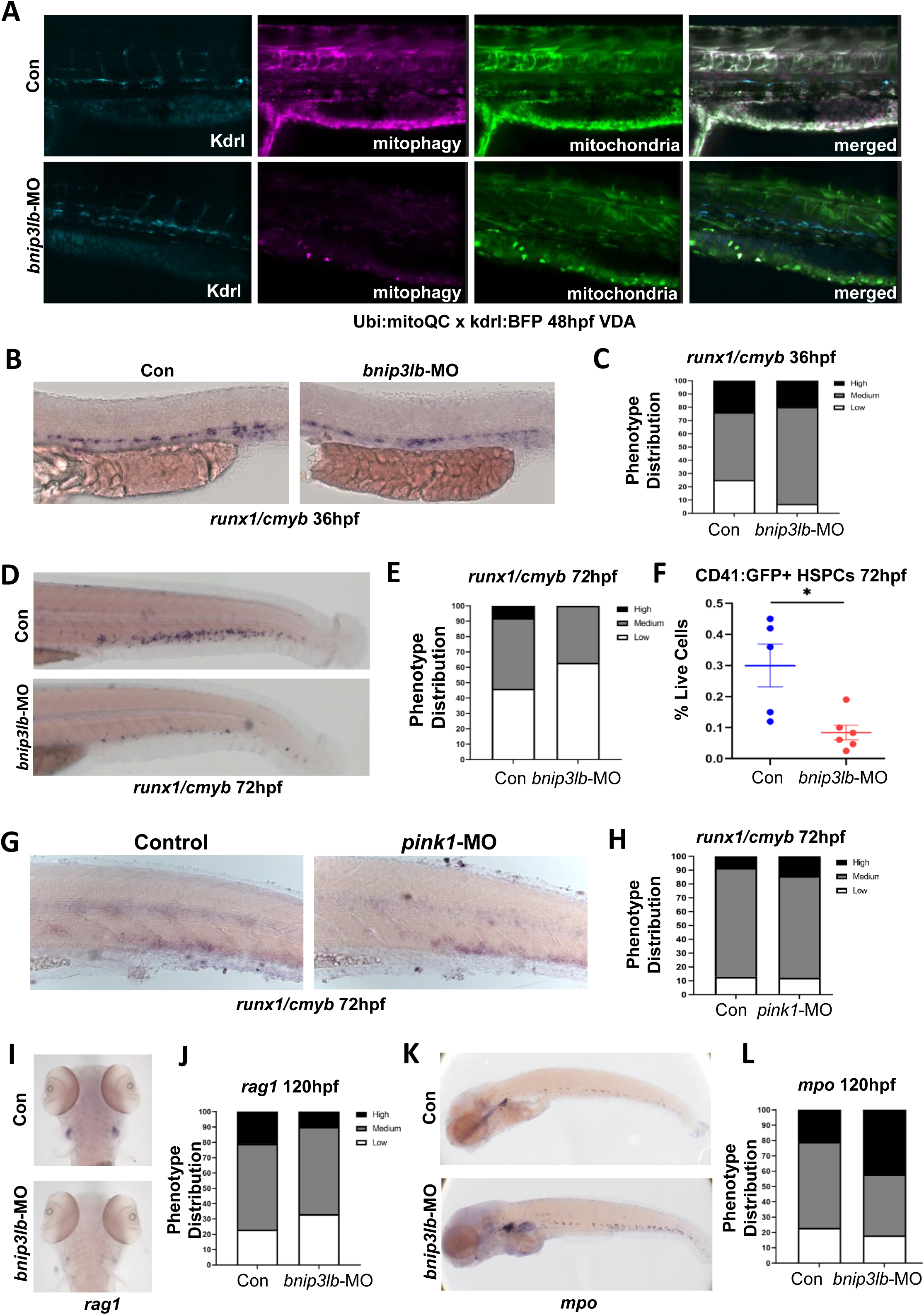
*Bnip3lb* knockdown reduces embryonic HSPC and lymphoid populations (See also Figure S4, S5) (A) Representative confocal image of mitophagy in the VDA at 48hpf, delineated by Kdrl expression, with and without *bnip3lb* knockdown, representative of 4 embryos (B): Representative WISH images of *runx1/cmyb* expression in the VDA at 36hpf in *bnip3lb* morphant embryos versus control (C): Phenotypic distribution of *runx1/cmyb* expression scored in embryos from (B) (D): Representative WISH image of *runx1/cmyb* expression in the CHT at 72hpf following *bnip3lb* knockdown (E): Phenotypic distribution of *runx1/cmyb* expression scored in embryos from (D) (F): Flow cytometry for CD41:EGFP+ HSPCs in *Tg(CD41:EGFP*) at 72hpf after *bnip3lb* knockdown, p=0.0110, Mean±SEM (n=8 embryos/clutch, 6 clutches) (G) Representative WISH images of *runx1/cmyb* expression in the CHT region at 72hpf after *pink1* knockdown (H) Phenotypic distribution of *runx1/cmyb* expression scored in embryos from (G) (I): Representative WISH images of *mpo* expression in the CHT and kidney marrow at 120hpf in *bnip3lb* morphants (J): Phenotypic distribution of *mpo* expression in KM scored in embryos from (F) (K): Representative WISH images of *rag1* in the thymus at 120hpf in *bnip3lb* morphants (L): Phenotypic distribution of *rag1* expression scored in embryos from (K)

**Figure 5:**
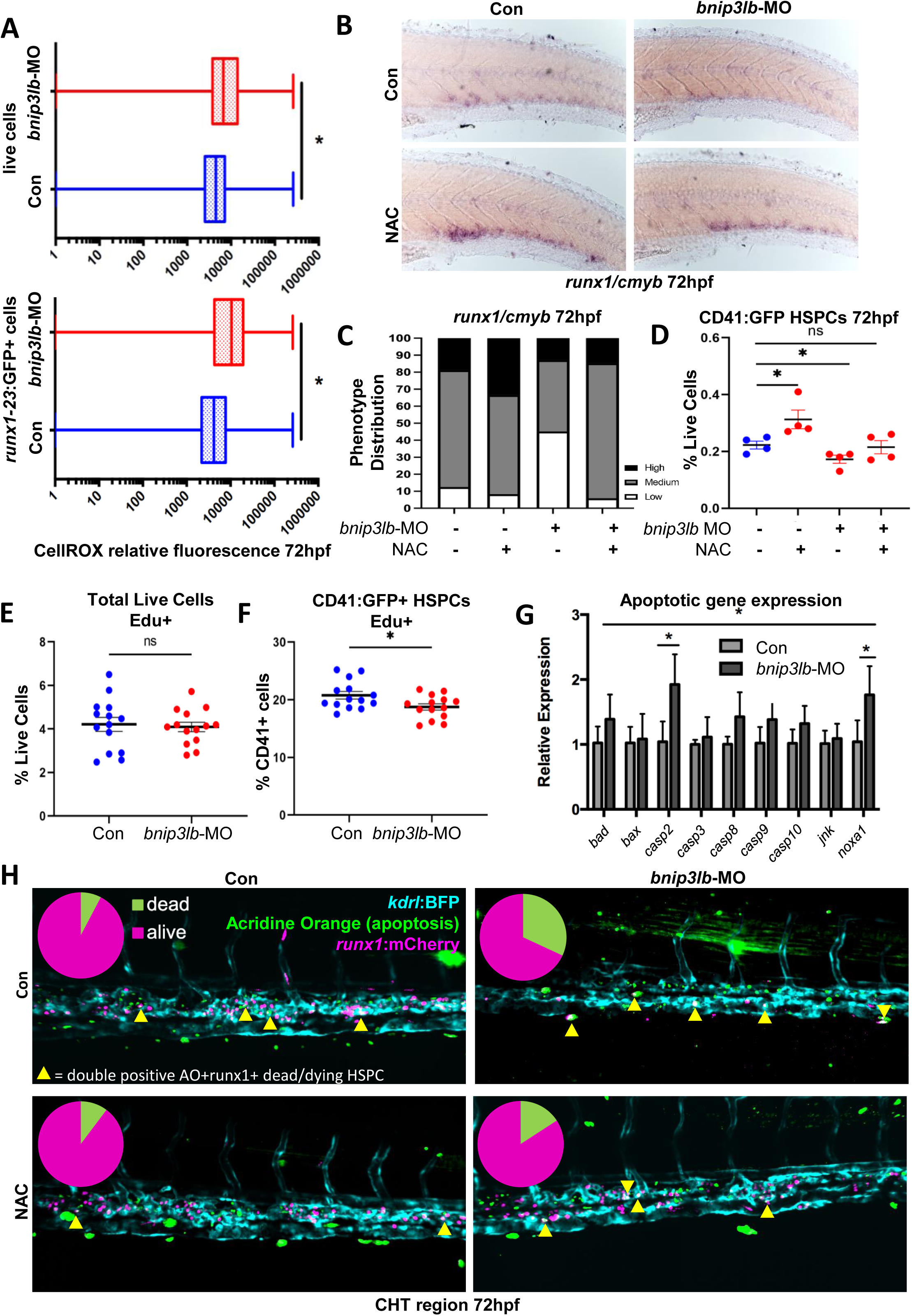
*Bnip3lb* protects embryonic HSPCs from ROS driven apoptosis. (A): ROS levels in 72hpf whole embryos (top) and *runx1* positive HSPCs (bottom) measured by flow cytometry for CellROX Orange^TM^ ROS-dye. Representative example of n=3, significance calculated by FlowJo K-S test (B): Representative WISH for *runx1/cmyb* expression at 72hpf in control and *bnip3lb* morphants (10uM NAC, 48–72hpf) (C): Phenotypic distribution of *runx1/cmyb* expression scored in embryos from (B) (D): Frequency of CD41:EGFP+ HSPCs by flow cytometry at 72hpf (10um NAC at 48hpf, Mean±SEM, n=8 embryos/clutch, 4 clutches) (D): EdU analysis in *bnip3lb* morphants by flow cytometry at 72hpf. (Mean±SEM, n=8embryos/clutch, 14 clutches, p=0.7609) (E): EdU analysis within the CD41:EGFP+ HSPCs fraction of *bnip3lb* morphants by flow cytometry at 72hpf (Mean±SEM, n=8 embryos/clutch, 14 clutches, p=0.0245). (F): qPCR for apoptotic markers at 72hpf after *bnip3lb* knockdown (significance for entire apoptotic panel: ANOVA p<0.001); individual significance for *casp2* (p=0.0197) and *noxa1* (p=0.0396) expression: t-test (Mean±SEM, n=12-15embryos/clutch, 3 clutches) (G): Confocal live imaging of HSPCs in CHT of *bnip3lb* morphants (10uM NAC tx:48-72hpf) at 72hpf. Kdrl+ vasculature (*blue*), acridine orange apoptosis (*green*), Runx1+HSPCs (*magenta*). Dual-labeled apoptotic Runx1+ cells indicated with yellow arrows. Pie charts show dead Runx1+ cells (*magenta*) as a proportion of the total, n=3/condition Chi square test, p=0.0443)

### *Bnip3lb* protects HSPCs from ROS driven apoptosis

Based on our primary observations and the impact of mitophagy in adult HSCs, we hypothesized that embryonic Bnip3lb-driven mitophagy is responsible for regulating ROS levels to promote survival and balance developmental lineage production with HSPC expansion to allow for establishment of the lifelong HSPC pool. To validate this conjecture, *bnip3lb* morphants were assessed by flow cytometry for CellROX, and ROS was found to be not only significantly elevated in the embryo as a whole, but to an even greater extent within the *Tg(Mmu.Runx1:EGFP)+* HSPC population (**Figure 5a**). To determine if increased ROS was responsible for the reduction of HSPCs in the CHT of *bnip3lb* morphants, we conducted an epistasis analysis with and without NAC treatment (48 to 72hpf). Knockdown of *bnip3lb* reduced HSPCs in the CHT at 72hpf, while NAC enhanced their appearance, as expected; furthermore, NAC-mediated reduction in ROS was sufficient to rescue HSPC gene expression after *bnip3lb* loss by WISH and by flow cytometry of CD41:GFP cells at 72hpf (**Figure 5b-d**).

We next aimed to clarify the mechanisms by which unregulated ROS impacts the embryonic HSPC population. ROS can promote or impede cell cycling, depending on the cellular context and dose^2^. Using a FACS-based EdU incorporation assay, we found that whereas the number of dividing cells in the whole embryo remained relatively constant, there was a significant reduction in the proportion of dividing CD41+ HSPC s in *bnip3lb* morphants (**Figure 5e, f**). While reductions in proliferation, combined with observed alterations in lineage commitment, may explain the impact of loss of mitophagy on HSPC maintenance in the CHT, high ROS can also drive apoptosis. Analysis of a panel of apoptotic genes at 72hpf by qPCR revealed a strongly significant response to *bnip3lb* knockdown overall, with significantly increased expression of the pro-apoptotic caspase *casp2*, and *noxa1*, a gene critical for mitochondria-mediated apoptosis^70^, in morphant embryos (**Figure 5g**). To confirm a potential role for cell death in the observed HSPC depletion present in *bnip3lb* morphants, we employed Acridine Orange, a marker for dead and dying cells. Assessment of the *Tg(Mmu.Runx1:NLS-mCherry)^cz2010^+* HSPC population revealed a pronounced enhancement in dual labeled cells in the CHT of *bnip3lb* morphants, indicative of increased apoptosis. Furthermore, this effect was rescued by reduction of ROS levels via NAC treatment, resulting in both less cell death and a greater number of Runx1+ cells overall (**Figure 5h**) in *bnip3lb* morphants.

Taken together, Bnip3lb*-*associated mitophagy was found to regulate the maintenance of newly derived HSPCs in the developing embryo by allowing them to proliferate while protecting them from ROS-induced apoptosis and/or premature lineage commitment.

### Promoting mitophagy enlarges the HSPC population and promotes lymphoid development

To further validate a beneficial functional role of mitophagy in the establishment and/or maintenance of the multi-potent HSPC population during embryonic development, we sought to understand whether stimulation of mitophagy could expand HSC production in vivo. To this end, we utilized an established ΔOTC construct, which employs a clinically-derived mutated form of mitochondrial matrix protein ornithine transcarbamylase to induce mitophagy via the misfolded protein response^64^. We engineered this gene under the heat-shock promoter to form an inducible *Tg(hsp70:ΔOTC)* mitophagy line.

Importantly, following heat-shock stimulation at 48hpf, dual transgenic *Tg(hsp70:ΔOTC;ubi:mitoQC)* embryos showed an increase in mitophagy reporter expression, particularly within the CHT (**Figure 6a**). In *Tg(hsp70:ΔOTC;CD41:EGFP)* embryos, ΔOTC induction (at 48hpf) drove a significant expansion in HSPC number at 72hpf by flow cytometry (**Figure 6b**); this positive impact of mitophagy stimulation was observed in the CHT as early as 48hpf by WISH for *runx1/cmyb* (**Figure 6c-f**). Furthermore, ΔOTC-stimulated mitophagy appeared to promote lymphoid lineage development, whereby a considerable increase in *rag1* occurred by 120hpf (**Figure 6g, h**), without affecting *mpo* expression as determined by WISH (**Supp. Figure 6a, b**).

**Figure 6:**
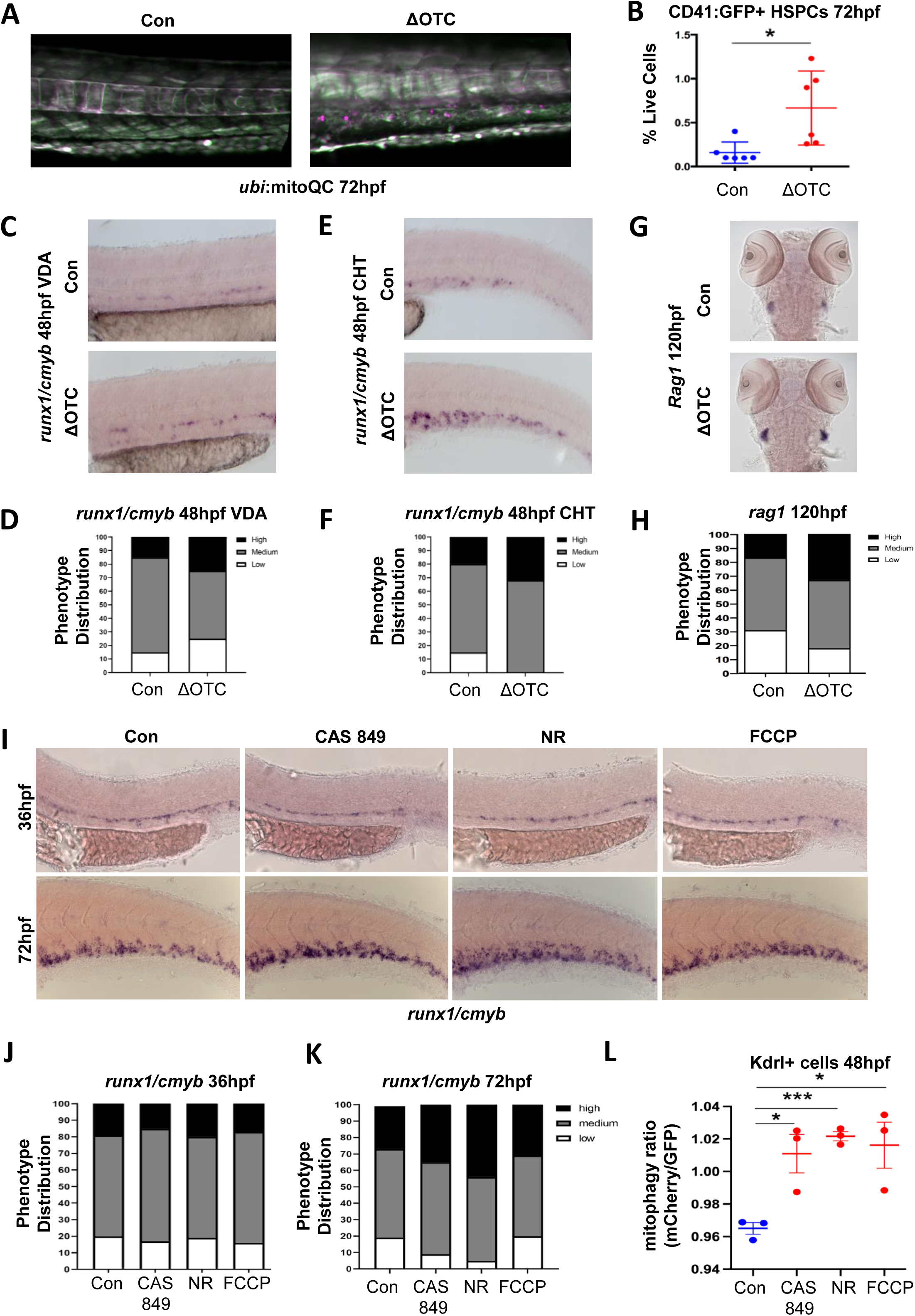
Induction of mitophagy enlarges the HSPC population and promotes lymphoid development (See also Figure S6) (A): Representative image of mitophagy in the CHT at 72hpf after *ΔOTC* gene expression (heat-shock at 38°C for 30mins at 48hpf). Healthy mitochondria (*green*), mitophagy (*magenta*). 5 larvae/condition. (B): Frequency of CD41:EGFP+ HSPCs by flow cytometry at 72hpf after induction of *ΔOTC* expression at 48hpf. (Mean±SEM, n=8embryos/clutch, 6 clutches, p=0.0176) (C): Expression of *runx1/cmyb* in VDA following *ΔOTC* induction at 24hpf assessed by WISH at 48hpf. (D): Phenotypic distribution of *runx1/cmyb* expression scored in embryos from (C) (E): Expression of *runx1/cmyb* in the CHT following *ΔOTC* induction at 24hpf assessed by WISH at 48hpf. (F): Phenotypic distribution of *runx1/cmyb* expression scored in embryos from (E) (G): Representative WISH images of thymic *rag1* at 120hpf, *ΔOTC* expressed by daily heat-shock from 48hpf (H): Phenotypic distribution of *rag1* expression scored in embryos from (I) (I): Representative WISH images of *runx1/cmyb* expression in embryos treated with CAS 849 (0.25uM), NR (10uM), or FCCP (0.01uM) from 24hpf-36hpf (*top row*) or 48hpf-72hpf (*bottom row*) to pharmacologically stimulate mitophagy. (J): Phenotypic distribution of *runx1/cmyb* expression scored in 36hpf embryos from (K) (K): Phenotypic distribution of *runx1/cmyb* expression scored in 72hpf embryos from (K) (L): Ratiometric analysis of mCherry/GFP (mitophagy) in Kdrl+ cells via confocal after treatment with CAS 849 (p=0.0204), FCCP (p=0.0247) or NR (p=0.0002). (Mean±SEM, n=12-15 fish/clutch, 3 clutches)

As a well-established issue with the derivation of clinical grade HSPCs from human iPSCs for therapeutic use is limited multi-potency and self-renewal capabilities, we next aimed to explore whether targeted stimulation of mitophagy might be functionally beneficial ex vivo. Chemical interventions are often simpler to perform and raise fewer regulatory concerns than genetic manipulations when attempting to translate animal model research into human cells intended for therapeutic use. Toward this end, we examined the impact of compound-mediated induction of mitophagy on HSPC development in zebrafish embryos. Several classes of chemicals have been previously noted to stimulate mitophagy in vitro and in vivo, including: carbonyl cyanide-trifluoromethoxyphenyl hydrazone (FCCP), which depolarizes mitochondrial membranes and engages the pink1-parkin pathway^71^; Nicotinamide riboside (NR), a NAD+ precursor which promotes transcriptional upregulation of mitophagy associated genes^72,73^; and AMPK activator CAS-84927-81-7 (CAS-849), which upregulates mitophagy via the mTOR pathway^74,75^.

Consistent with our spatio-temporal expression data for *bnip3lb*, we saw little effect of mitophagy stimulation on specification of HSPCs in the VDA at 36hpf. In contrast, *runx1/cmyb* expression notably increased in the CHT at 72hpf by WISH following NR and CAS-849 treatment (**Figure 6i-k**). This could be observed as early as 48hpf, as HSPCs migrate from the VDA to CHT (**Supp. Figure 6c, d**), suggesting compound-mediated stimulation of mitophagy may be a viable avenue to promote HSPC expansion following EHT. Interestingly, while all three compounds stimulate mitophagy in vivo at 48hpf (**Figure 6l**), the impact of FCCP on HSPCs was minimal, perhaps reflective of its mechanism of action via mitochondrial membrane damage, leading to ROS-associated cellular toxicity and/or reliance on the Pink1-Parkin pathway.

### NR-driven mitophagy promotes colony forming potential in hiPSC derived hematopoietic progenitors

In addition to being the most efficient of the tested compounds in enhancing mitophagy in zebrafish embryos, NR is well tolerated in vivo and in vitro^72,73^. In line with our WISH observations, NR treatment significantly increased the CD41+ HSPC population of the CHT by flow cytometry (**Figure 7a**). Furthermore, sustained exposure continued to positively influence HSPC number during colonization of their terminal niche, the larval KM (**Figure 7b, c**), and significantly increased the *Tg(Rag2:GFP)* lymphoid population in the thymus (**Figure 7d**). We therefore explored use of NR-mediated stimulation of mitophagy for our translational studies.

**Figure 7:**
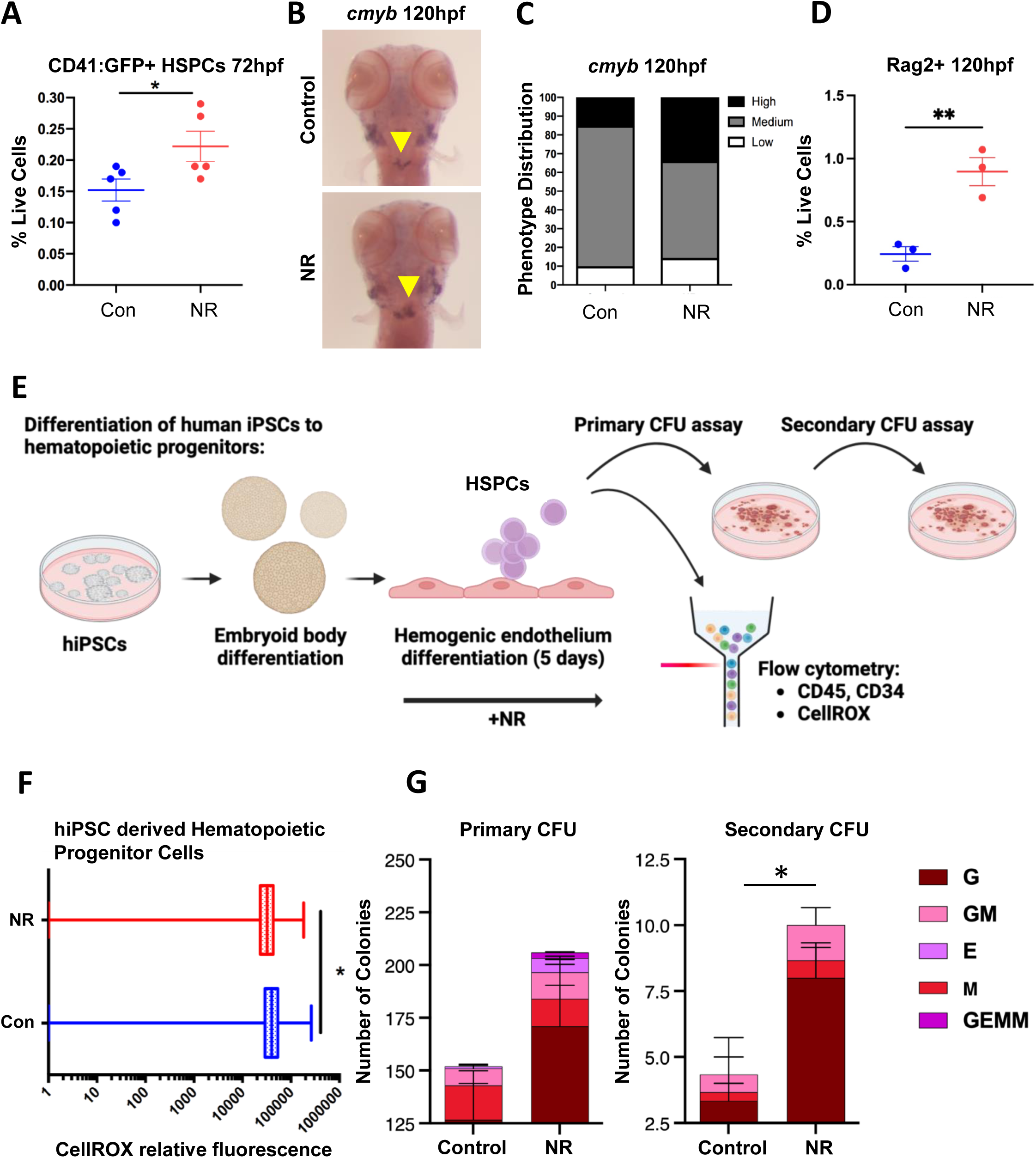
NR-driven mitophagy promotes colony forming potential in hiPSC derived hematopoietic progenitors (See also Figure S7) (A): Frequency of CD41:EGFP+ HSPCs by flow cytometry at 72hpf (10uM NR; 48hpf-72hpf). Mean±SEM, n=8embryos/clutch, 6 clutches, p=0.0477) (B): Representative WISH image of *cmyb* expression in the KM (*yellow arrows*) at 120hpf (10uM NR; 48hpf-120hpf) (C): Phenotypic distribution of *runx1/cmyb* expression scored in embryos from (B) (D): Frequency of Rag2+ lymphoid progenitor cells by flow cytometry at 120hpf (10uM NR 48hpf-120hpf). Mean ± SEM, n=8embryos/clutch, 4 clutches p=0.0178) (E): Schematic of hiPSC differentiation protocol: hiPSCs undergo step-wise endothelial commitment, then EHT. Non-adherent cells analyzed by flow cytometry for CellROX, CD45 and CD34. FACS-isolated CD34+ HSPCs were utilized in serial-replating CFU assays. NR treatment (1mM) conducted at EHT stage. (G): CellROX Orange^TM^ ROS-dye indicates a decrease in ROS levels in NR-treated (1mM) CD34+ HSPCs; significance calculated by FlowJo K-S test (H): CFU assay after NR treatment showing distribution of colony-types scored at after primary (left, day 14 post plating) and secondary (right, serial) culture compared to control (n = 3)

Human iPSCs were differentiated toward hematopoietic fate using a modified Keller protocol^76^ and treated with NR (1mM) during the EHT stage (**Figure 7e)**. Comparable to the lack of impact on HSPC specification in the zebrafish VDA, no significant differences were found in the numbers or proportions of CD34+CD45+ multipotent hematopoietic progenitors initially generated (**Supp. Figure 7**) at 5 days of culture. Intriguingly, however, NR was noted to reduce intracellular ROS levels in all cultured cells as measured by CellROX^TM^ at this timepoint (**Figure 7f**). Following flow cytometric profiling, control and NR-treated cells were collected for assessment of hematopoietic colony forming potential using standard methylcellulose colony forming (CFU) assays. At day 14 of the primary CFU assay, there was a dramatic increase in all colony types scored following prior NR-stimulation compared to controls; this included the presence of multipotent CFU-GEMM colonies, which were not observed in control samples (**Figure 7g**). To assess potential long-term impact on HSPC production and/or function, cells were harvested and serially replated into secondary CFU culture: at 28 days from initial harvest, increases in all colony types represented were retained in NR-treated samples (**Figure 7g**), indicating that induction of mitophagy promotes maintenance of de novo-derived human multi-potent hematopoietic progenitors in vitro.

Altogether, our data indicates that activation of developmentally programmed mitophagy via Bnip3lb function is an essential component of HSPC development in the vertebrate embryo, which regulates ROS levels to allow for the expansion, controlled differentiation, and survival of the stem cell pool. Furthermore, these findings appear to have direct therapeutic implications for enhancing the maintenance and functional capacity HSPCs derived de novo in vitro from human iPSCs.

## DISCUSSION

HSC transplantation is the gold standard of treatment for many otherwise incurable hematopoietic malignancies. As such, generation of patient-specific HSPCs from hiPSCs could alleviate major issues surrounding matched donor availability and GVHD. However, standard in vitro differentiation protocols do not allow for formation or maintenance of hiPSC-derived HSCs with long-term multi potency and self-renewal capabilities at a scale sufficient for clinical use ^77,78^. Improvement of ex vivo production protocols, therefore, requires better understanding of the regulatory interactions governing in vivo HSC development and maturation. Whereas most developmental studies have focused on initial specification of HSPCs and the process of EHT, analysis of CHT/fetal liver stages of hematopoiesis provides an opportunity to model the unique developmental phenomena whereby HSPCs can divide many times while maintaining “stemness”. The work detailed here regarding developmentally programmed mitophagy in HSPCs expands our understanding of the regulatory interactions at play in these secondary “expansion” hematopoietic niches. The primary observations presented support conjectures from our lab, and others, that precise regulation of metabolic and inflammatory state has continual influence on HSCs throughout developmental hematopoiesis, affecting both the production and long-term function of the nascent HSC pool^49,79,80^.

The impact of ROS during developmental hematopoiesis was of particular interest to us given it potently enhances HSPC specification in the VDA^52^, but is known to damage or destroy the HSPC population in the adult BM^10^. This implied the presence of a temporal or spatially restricted ‘switch’ in the role of ROS during development, and necessity for a means of timed ROS regulation. In this study, we found excess ROS, produced during the shift to embryonic OX/PHOS metabolism, can become toxic or detrimental to newly specified HSPCs immediately after EHT. This directly coincides with initiation of *bnip3lb* gene expression, which is precisely coordinated as cells transition from an HEC to HSC transcriptional profile. Indeed, induction of *bnip3lb*-regulated mitophagy can be observed in the presumptive HSPC population as they progress through EHT in the VDA, and is maintained in CD41+ cells in the CHT. Importantly, our *bnip3lb* knockdown studies indicate that mitophagy is essential for reducing the amount of ROS within the HSPC population in the CHT, which protects them from apoptosis, while allowing both proliferation and differentiation to progress appropriately. The importance of maintaining this ROS balance is emphasized in a recent report by Wattrus et al.^81^, which showed HSPCs in the CHT with excessive ROS were engulfed by macrophages to maintain the quality of the HSPC pool; however, the mechanism by which ROS levels were balanced to protect the majority of HSPCs from phagocytosis was not previously characterized. Mechanisms of ROS regulation have also been explored in murine models: ablation of Fip200^82^, an essential component of the autophagic isolation membrane, revealed similar loss in fetal HSCs as reported here, suggesting autophagy is likely employed across species to control metabolic state and maintain stemness in newly specified HSCs.

While we showed Bnip3lb induction is responsive to environmental cues in the niche such as ROS and hypoxia, its expression does not appear to be uniquely dependent on these factors. While we were unable to define a singular regulatory pathway responsible for onset of *bnip3lb* expression in the VDA, we speculate it may be coordinately or redundantly initiated as part of, or in response to, the implementation of a larger developmental program during the process of EHT. Such developmentally programmed regulatory control of Bnip3lb expression is consistent with literature in both related and disparate cell types, including the maturation of red blood cells and lens fibers in the developing eye^42–47^. We note that our data showed no relevance for Pink1-Parkin pathway activation in response to oxidative or other stressors in the CHT; this is in contrast to adult HSCs, which use Pink1-Parkin, but not Bnip3l/Nix^11,19,34^, further supportive of our presumption of directed employment of a mitophagy program specific to these developmental stages. It is important to state, however, that our transcriptional profiling does suggest de novo specified HSPCs exhibit a classical “stress-response” gene signature, perhaps indicative of the overlapping regulatory routes or gene networks that can converge to ensure induction and/or maintenance of Bnip3lb-associated mitophagy. Aligning with that conjecture, a recent paper by Liu et al.^83^ showed Atg5-dependent autophagy is essential for EHT, highlighting the importance of multiple autophagic pathways in embryonic HSC production. Developmentally programmed mitophagy is an under-studied phenomenon in general, and our discoveries contribute to a mounting body of evidence suggesting its use in embryonic tissue specification, and perhaps stem and progenitor cell biology in general, is more widespread than currently reported^33^.

Finally, our data imply that attempts to generate or maintain HSCs *in vitro* should simultaneously seek to recapitulate induction of mitophagy, to aid in preserving full self-renewal and multi-potency of newly specified progenitors. Toward that aim, we identified several effective small molecule chemical mitophagy activators that enhanced HSC numbers and lymphoid progenitor production *in vivo,* selecting the NAD+ precursor NR for further in vitro translational studies given its safety profile^84^. NR exposure during human iPSC hematopoietic differentiation significantly improved hematopoietic colony forming potential, particularly in the context of serial replating, reflective of improved HSPC maintenance. Of note, NR regulates cellular redox via multiple pathways and is not specific to mitophagy alone, so additional work will be needed to distinguish the isolated and combinatorial impact of each potential arm of its regulatory mechanism. Nevertheless, our findings contribute to our burgeoning mechanistic understanding of regulatory interplay in secondary “expansion” niches by highlighting the essential role of developmentally programmed, non-mitochondrial-damage-dependent mitophagy in HSPC development, and the fine spatio-temporal requirement for Bnip3lb/Nix in particular. Further, this work highlights a potential adjuvant path towards in vitro generation and/or expansion of multi-potent self-renewing HSPCs for transplant therapy, and therefore improved clinical outcomes for patients in need.

## STAR METHODS

### Resource availability

#### Lead Contact

Requests for resources should be directed to and will be fulfilled by the lead contact, Trista E. North.

#### Materials availability

Materials generated in this study are available upon request.

#### Data and Code availability

scRNAseq data will be deposited at GEO and will be publicly available as of the date of publication. Accession numbers will be listed in the [key resources table](https://www.ncbi.nlm.nih.gov/pmc/articles/PMC9386785/table/T1/). Any additional information required to reanalyze the data reported in this paper is available from the lead contact upon request.

## Experimental model details

### Zebrafish Husbandry

Zebrafish were utilized in accordance with protocols approved by Boston Children’s Hospital and Beth Israel Deaconess Medical Center Institutional Animal Care and Use Committees (Protocols: 0002155 and 045-2021). Adults were maintained in a standard circulating water facility on a live brine shrimp diet supplemented with Gemma 300 pellets (Skretting). Wild-type strains used were *AB* or *TU*; transgenic and/or mutant lines utilized are detailed in the Key Resources Table. Mutants were genotyped by PCR on tail-clip samples (see Key Resources Table for primers). After screening for developmental abnormalities, healthy embryos were maintained in E3 fish medium in a 28°C incubator until use.

### hiPSC Culture

hiPSCs were provided by the Boston Children’s Hospital hESC (human embryonic stem cell) Core Facility and cultured in Stemflex medium (ThermoFisher Scientific) in 37°C, 20% O_2_, 5% CO_2_ incubators. Karyotyping and mycoplasma testing was performed to maintain quality control.

## Method details

### Chemical treatments

Embryos were exposed to compounds (see Key Resources Table) dissolved in E3 (fish) water at the concentrations and timepoints shown in Methods Table S1, with treatment of equivalent percentage of vehicle (DMSO) utilized for sibling controls.

### Flow Cytometry and Cell Sorting

Embryos (pools of 8 embryos/sample) were dissociated in 500μL Liberase^TM^ solution (75μg/mL in 1xPBS/1mM EDTA) at 34°C for 45 minutes. Samples were dissociated by pipetting and filtered. SYTOX Red viability stain was added to a concentration of 5nm. Flow cytometry was performed on an LSRFortessa (BD) and analyzed by FlowJo software. For ROS detection, CellROX orange dye (1:1000 dilution) was added to samples 30 mins prior to analysis and measured using 561nmn laser and PE filter. For cell sorting, embryos were bisected along the transverse plane of the trunk at the point where the yolk sac narrows, and tail sections dissociated as above. Sorting for live cells was performed on a FACS Aria (BD).

#### In situ hybridization

Whole mount in situ hybridization (WISH) was performed as described (http://zfin.org/ZFIN/Methods/ThisseProtocol.html). For each condition, three clutches of **≥**20 stage matched embryos were fixed in 4% paraformaldehyde. Fixed embryos were treated with 0.8%KOH/0.9%H_2_O_2_ to remove pigment, then dehydrated in methanol. Once rehydrated in PBS 0.1% Tween (PBST), embryos were permeabilized using 10μg/mL Proteinase K, re-fixed in 4% paraformaldehyde, then washed again in PBST. Hybridization with digoxigenin-UTP-labeled riboprobes was performed in hybridization buffer containing 50% formamide at 70°C overnight. Published probes are shown in the Key Resources Table; the *bnip3lb* probe was made from cDNA of the coding sequence using primers shown in the Key Resources Table. Embryos were washed in sodium citrate buffer/0.1% Tween and incubated overnight with Anti-dioxigenin-AP, Fab fragments. After washing in PBST, antibodies were visualized by incubation in Tris buffer (pH 8.5) with BCIP/NBT. Intensity of staining was scored qualitatively, with each embryo assessed as high, medium or low relative to the median intensity of wild-type controls from the same clutch. The scoring distribution of each sample set was averaged across three independent clutches, and control and treatment groups compared. Representative images were captured using a Zeiss Axio Imager A1/Axio Cam MRC and Axiovision LE software. Fluorescent WISH was performed as described^85^, by a procedure comparable to conventional WISH, but utilizing Anti-dioxigenin-POD, Fab fragments used to stain the *bnip3lb* probe, Anti-Fluorescein-POD, Fab fragments for the *runx1* probe, and Rabbit anti-GFP TP401 and Goat anti-Rabbit IgG Alexa fluor 647 to detect Kdrl:GFP. Fluorescent WISH images were captured using confocal microscopy as described below.

#### Confocal Imaging

Confocal imaging was performed on a Ti2 (Nikon) scope using an iXon Life 888 EMCCD (ANDOR) camera and analyzed using Imaris software (Oxford Instruments) and Python. Embryos were treated with 0.003% PTU at 24hpf to inhibit pigmentation. For live imaging, embryos were mounted on glass-bottom dishes in 1% low-melt point agarose in E3 media containing 0.2 mg/ml Tricaine. For acridine orange experiments, 72hpf embryos were exposed to 10 µg/mL of acridine orange in E3 media for 1 hour while protected from light, then washed in E3 three times before live imaging. Images were de-noised with the bm3d algorithm^86^ where indicated and are otherwise unaltered. HyPer relative fluorescence was quantified by masking on mCherry and taking the average values of pixels in the HyPer channel. Ratiometric analysis of the mitophagy reporter was performed as described^63^: a ratio of the mCherry/GFP values for each pixel was taken, and the results were averaged to give the final value reported, statistical significance determined with students t-test.

#### qPCR

RNA was extracted from mechanically homogenized pooled embryos (n = 20/condition) using RNAEasy Micro Kits according to the manufacturer’s instructions. Reverse transcription of 1ug of RNA was performed using a High-Capacity cDNA Reverse Transcription Kit according to the manufacturer’s instructions. SYBR Green PCR Master Mix was used for qPCR with an ABI7900 (Applied Biosystems) according to the manufacturer’s instructions. Relative expression was calculated using the ΔΔCT method^87^ See table S2 for qPCR primers.

#### scRNASeq

Two replicate samples of pooled dissected tail preps of 48hpf Kdrl+ embryos (300 embryos/condition) were isolated by FACS as described above. Samples were processed on a Chromium Next GEM Single Cell 3’ GEM, Library & Gel Bead Kit v3.1 and sequenced on a Novaseq 6000 system (Illumina). Data was processed using the Seurat^88^ and clusterProfiler^89^ packages.

#### Morphants

To create morphants, 1-3nl of a given morpholino oligonucleotide was injected into the yolk ball of embryos at the single cell stage at concentrations shown in the Key Resources Table, using a microinjector (Narishige) and a pulled glass needle. Uninjected sibling embryos were used as controls.

#### Transgenesis

*Tg(CD41:mitoQC)* was generated using tol2 transgenesis as described by Wrighton et al ^5^. Briefly, gateway cloning using LR Clonase II was used to make a pTol2-*Cd41:mitoQC, cryaa:Cerulean* vector, from p5E-CD41, pME-mitoQC, p3E-polyA and pDestTol2pA2 according to the manufacturer’s protocol. This vector along with *tol2* transposase mRNA, made using the mMessage mMachine SP6 *in vitro* transcription kit, was injected directly into fertilized embryos at the one cell stage. F0 embryos were screened for cerulean fluorescent lenses and outcrossed as adults. Potential founders were screened for brightness, stable integration and copy number. *Tg(hsp70:ΔOTC)* was generated as above using p5E-hsp70, pME-ΔOTC, p3E-polyA and pDestTol2pA2.

#### EdU Assay

Larvae were exposed to 500μM EdU in E3 water (10% DMSO) for 1 hour at 4°C. Pooled larvae were dissociated and filtered as above, then cells were fixed, permeabilized and stained as per Click-iT™ EdU Alexa Fluor™ 647 Flow Cytometry Assay Kit. Flow cytometric data were collected on LSRFortessa (BD) and analyzed using FlowJo. *Tg(CD41:EGFP)^+^*.

#### hiPSC differentiation

Human 1157.2 iPSCs^49^ were maintained on Matrigel matrix using Stemflex media and differentiated into CD34^+^45^+^ HSPCs cells using a previously published protocol^76^. Briefly, iPSC colonies were scraped, then cultured in ultra-low attachment plates with aggregation media containing BMP4 (10ng/ml, day 0-2), bFGF (5ng/ml, day 1-2), CHIR99021 (3μM, day 2), and SB431542 (6μM, day 2), VEGF (15ng/ml, day 3-8), bFGF (5ng/ml, day3-8), SCF (50ng/ml, day 6-8), EPO (day 6-8), IL-6 (day 6-8), IL-11 (day 6-8), and IGF-1 (day 6-8) to allow the formation of embryoid bodies (EBs). On day 8, EBs were dissociated using an EB Dissociation kit (Miltenyi), and CD34+ hemogenic endothelium (HE) cells isolated by magnetic-activated cell sorting (MACS) using the human CD34 Microbead Kit (Miltenyi). CD34^+^ HE cells were treated with DMSO or 1mM Nicotinamide Riboside (NR) and cultured for 5 days in StemPro-34 media (Invitrogen) supplemented with BMP4, bFGF, IL3, IL6, IL11, IGF-1, VEGF, SCF, EPO, TPO, Flt-3L, and SHH according to standard protocols^76^ to allow for the formation of non-adherent CD34^+^45^+^ HSPCs.

### Hematopoietic Colony Forming Unit (CFU) assay

CFU assays were performed using MethoCult H4636 methylcellulose media following manufacturer’s instructions. Colonies were counted after 14 days of culture; cells were then collected, washed with PBS and replated at 30k cells/well for secondary CFU assays.

### Quantification and Statistical Analysis

Data analyses were performed using GraphPad Prism and Excel. Statistical significance was determined by two-tailed Student’s T-test, unless otherwise indicated. Error bars indicate mean and SD, unless otherwise indicated.

### Key resources table

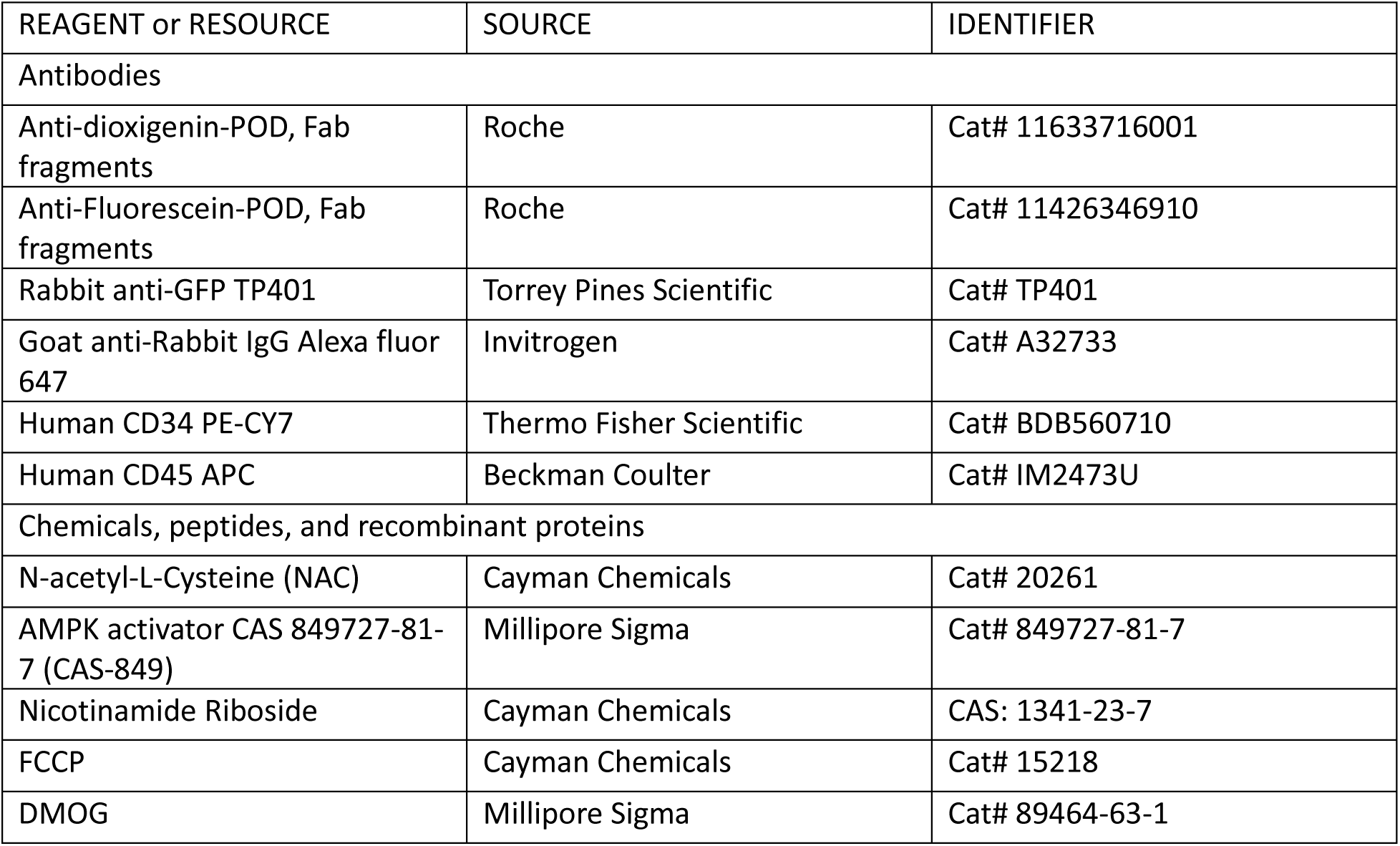
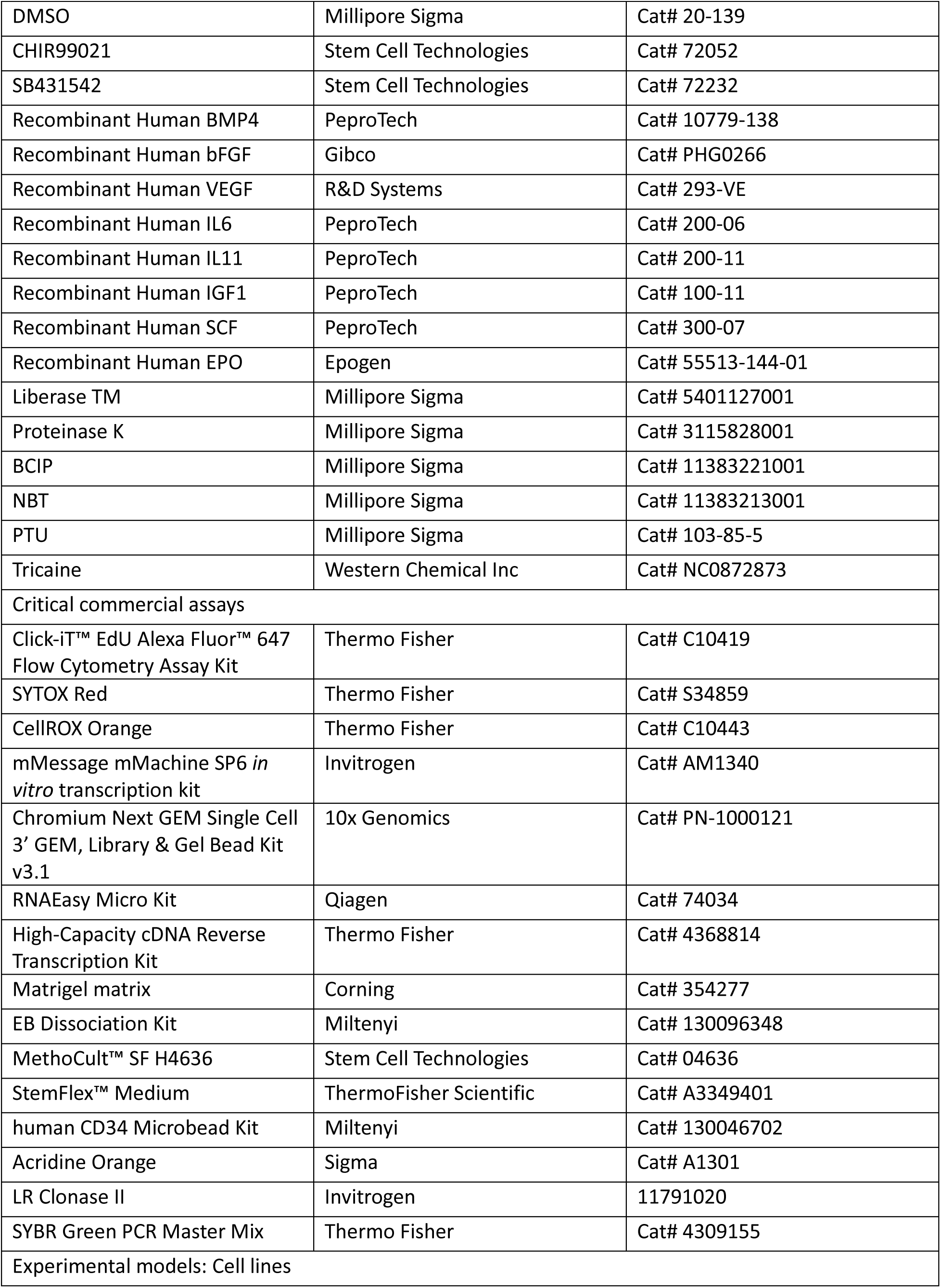
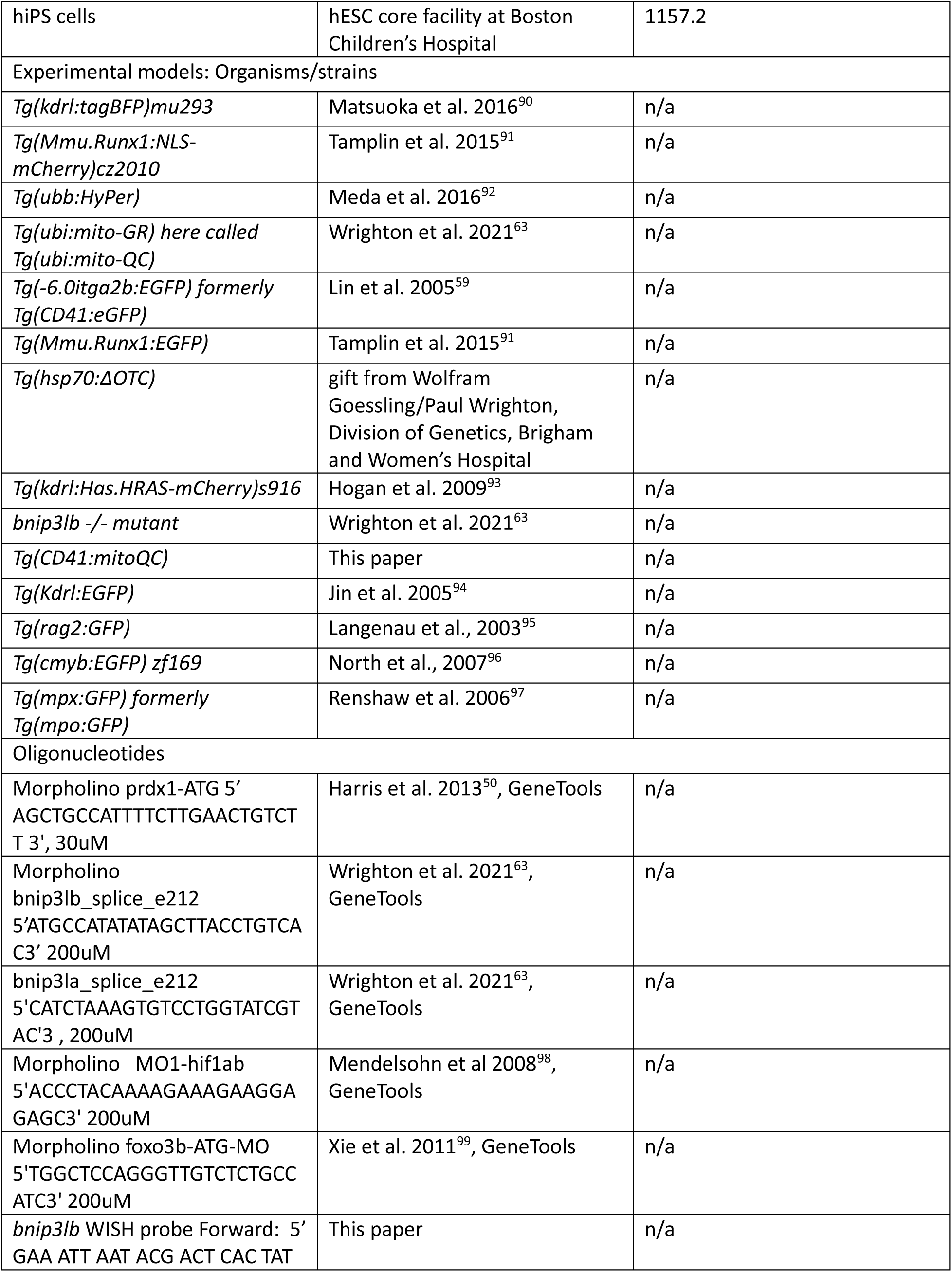
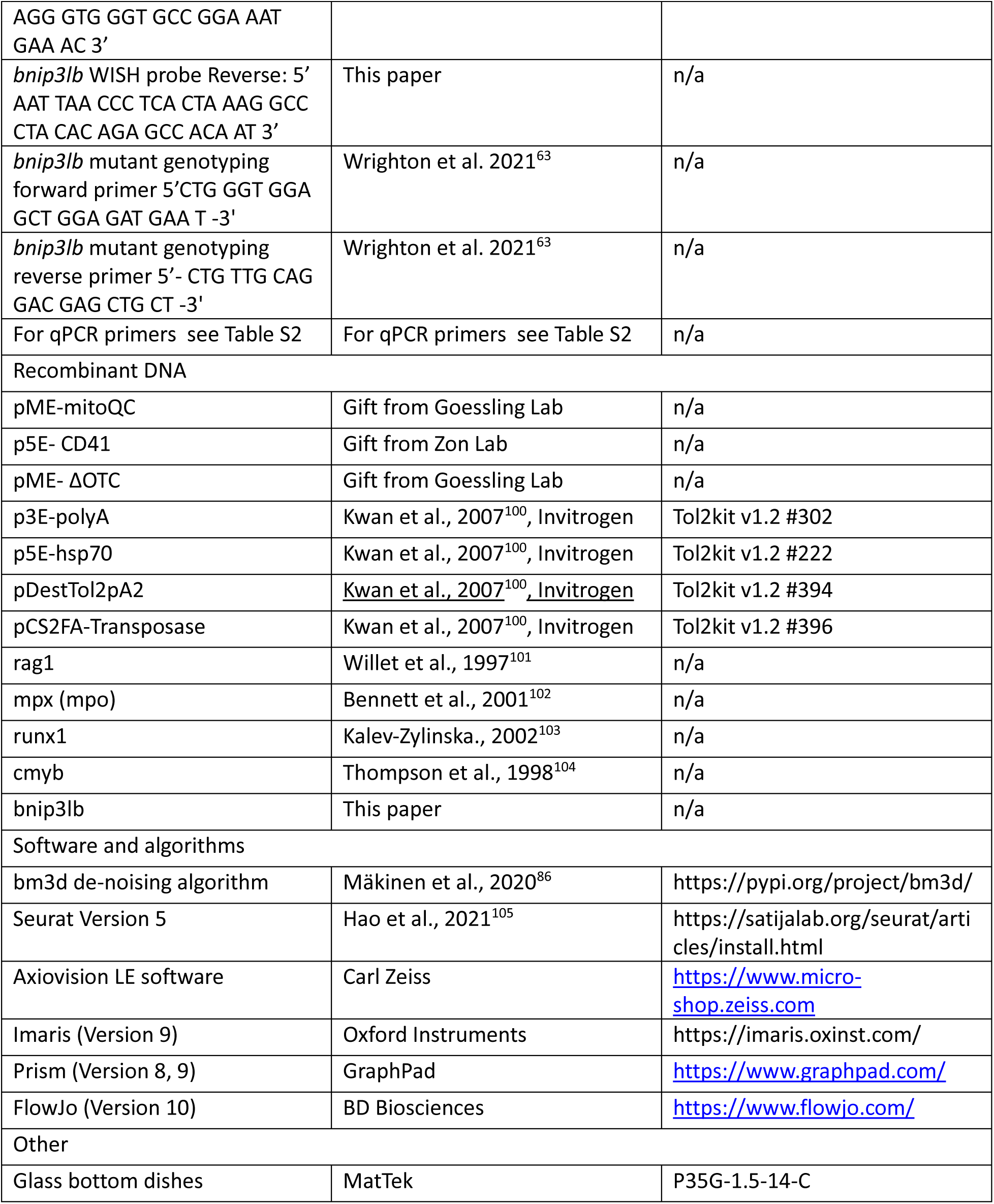

## Acknowledgements

We thank Boston Children’s Hospital (BCH) Flow Cytometry Core, BCH hESC Core Facility, Harvard University Single Cell Core, Harvard Chan Bioinformatics Core and Harvard University Biopolymers Facility for assistance. We are grateful to W Goessling and LI Zon for sharing materials and strains. We also thank E Molnar, J Frame, M Hachimi and TL Long for assistance and advice. This study was supported by NIH grants: 1R01HL152636 (TEN), 1R01HL154580 (TEN), 1RC2DK120535 (GQD/TEN).

## Author Contributions

Conceptualization: EM, TEN, GQD and WG; Methodology: EM, TEN; Validation: EM, MRL, VMT, MTW; Formal Analysis: EM, MAN, IMO; Investigation: EM, MTW, MRL, RJ, ZCL, PJW, VMT, WWS, S-EL, EDQ; Resources: TEN, WG, GQD.; Data Curation: EM; Writing: EM, TEN, MTW, MRL; Visualization: EM, MTW; Supervision: T.E.N. G.Q.D., W.G.; Project administration: E.M., T.E.N.; Funding: TEN, GQD.

## Supplemental Figures

**Supplemental Figure 1:**
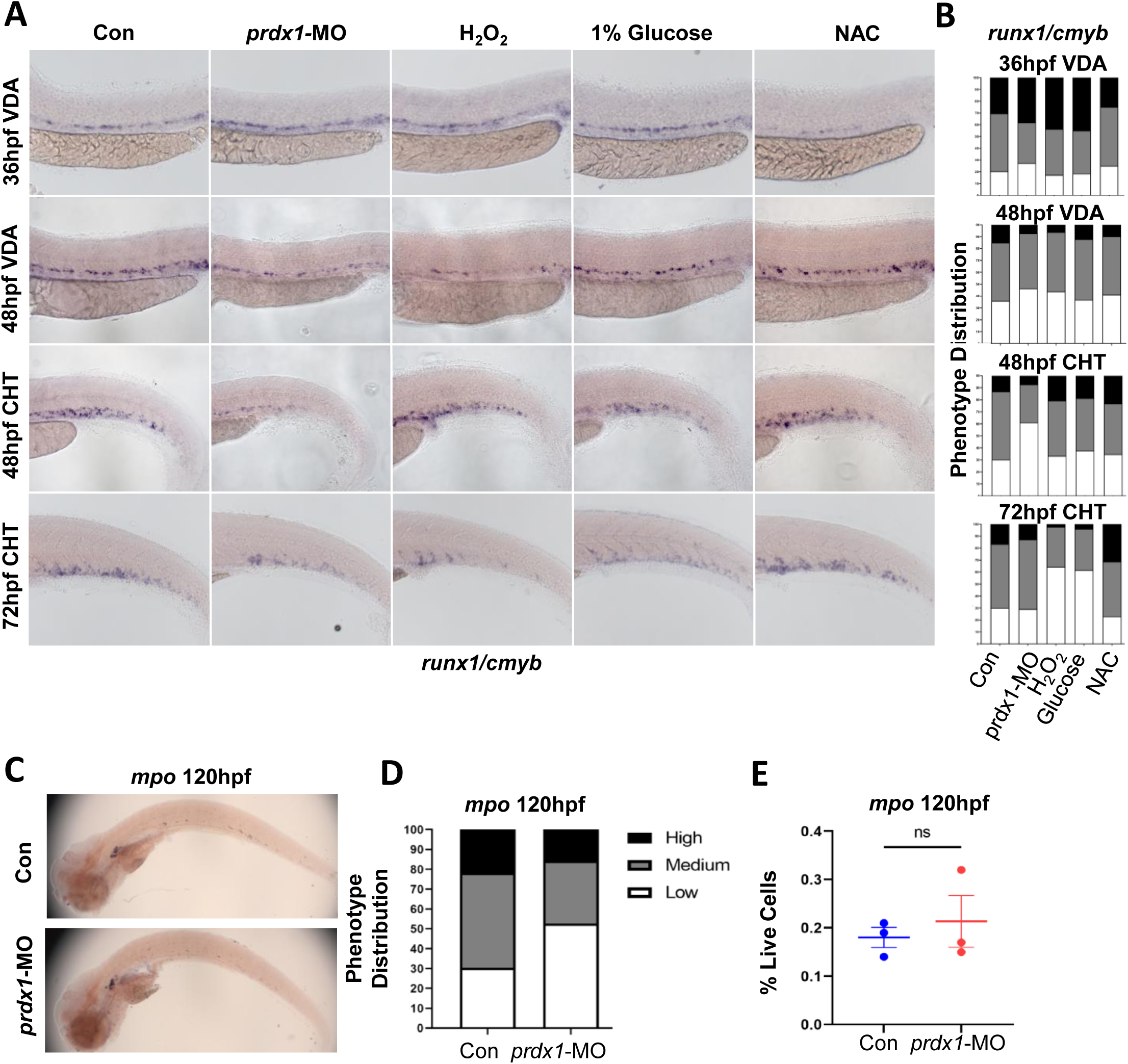
ROS regulates HSPC development differently in the VDA vs the CHT. (A): Representative image showing expression of *runx1/cmyb* in embryos with or without morpholino (MO)-mediated *prdx1* knockdown or H_2_O_2_, 1% Glucose, NAC treatment assessed by WISH at 36hpf, 48hpf and 72hpf (B): Phenotypic distribution plot of *runx1/cmyb* expression scored in embryos from (A) (n = 20 per clutch, 3 clutches) (C): Representative WISH images of *mpo* at 120hpf, after knockdown of *prdx1* with a MO (D): Phenotypic distribution plot of *mpo* expression scored in embryos from (C) (n = 20 per clutch, 3 clutches) (E): The frequency of *mpo*+ cells by FC at 120hpf. Mean ± SEM, n=8 embryos per clutch, 3 clutches

**Supplemental Figure 2:**
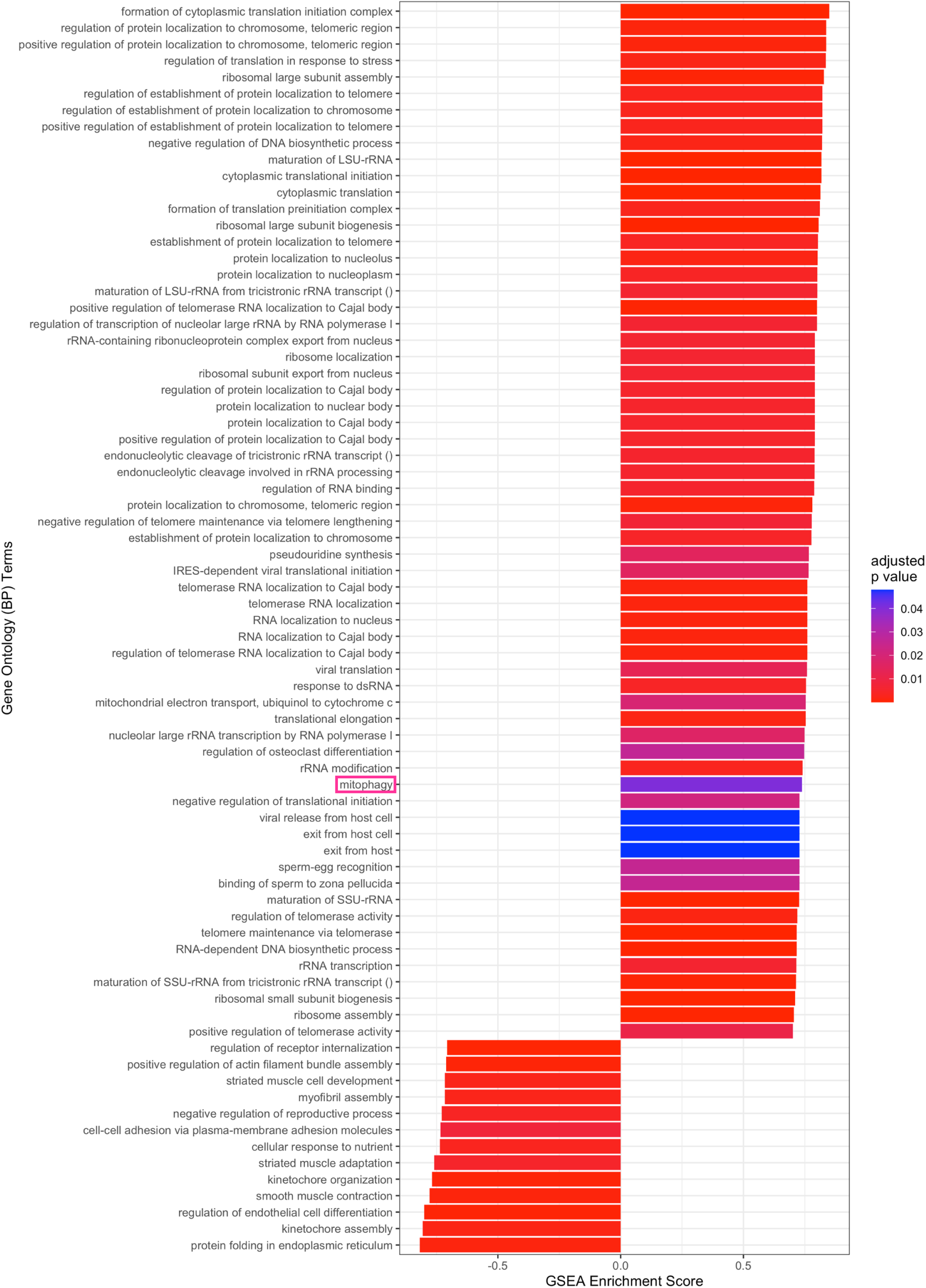
GSEA analysis shows mitophagy process enriched in HSPC cluster (Related to Figure 2) GSEA analysis showing enrichment scores of Gene Ontology:Biological Processes terms enriched in HSPC cluster compared to hemogenic endothelium, generated from deseq2 analysis.

**Supplemental Figure 3:**
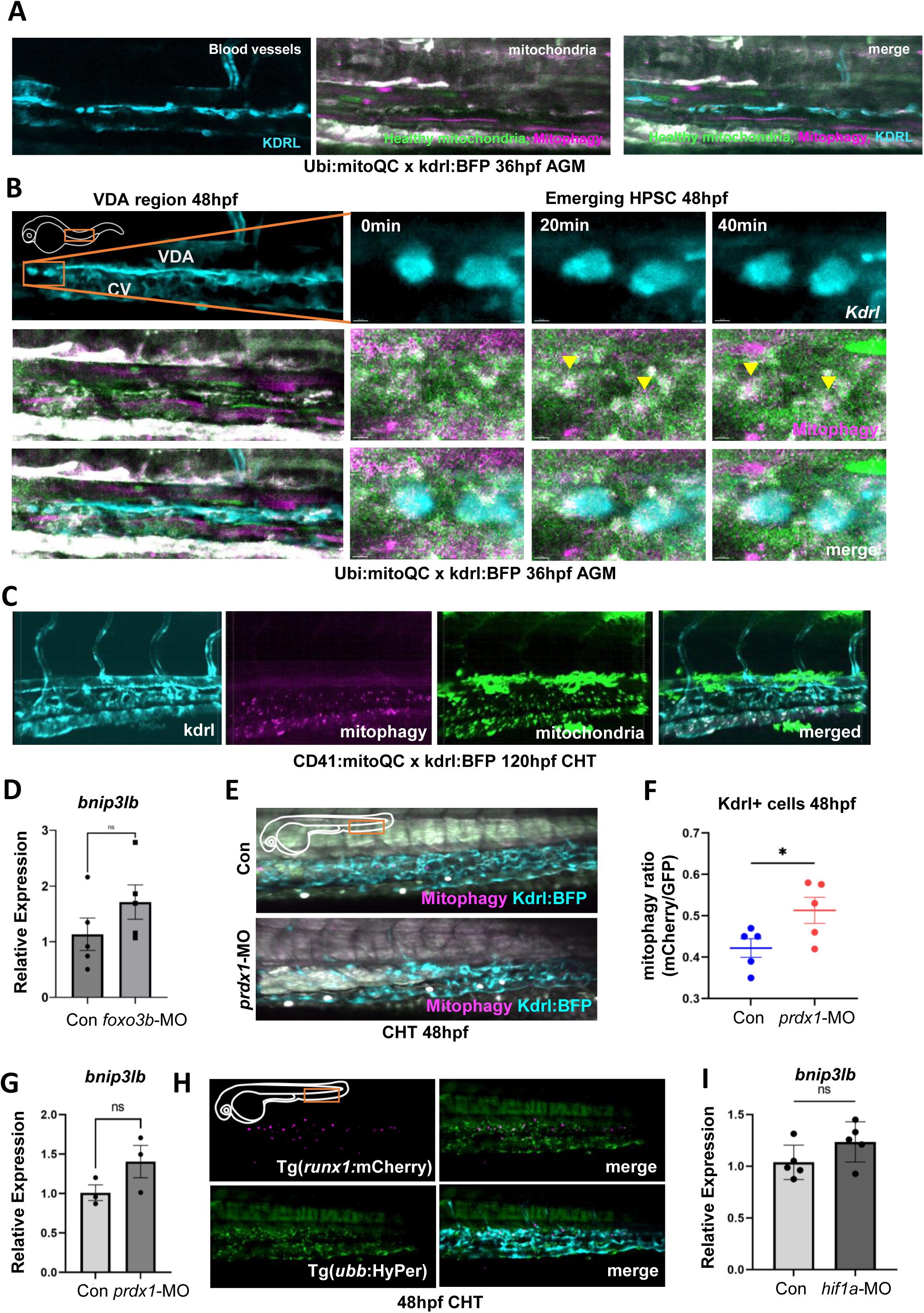
ROS induces an increase in mitophagy within the CHT (Related to Figure 3) (A): Timelapse video of confocal images shown in figure 3b showing mitophagy occurring in the VDA region at 48hpf. Healthy mitochondria (green), mitophagy (magenta), *kdrl* (blue). (B): Additional confocal timelapse images of two more budding cells with mitophagy occurring in the AGM at 48hpf. Healthy mitochondria (green), mitophagy (magenta), *kdrl* (blue). (C): Confocal image of *Tg(CD41:mitoQC)* larva showing mitophagy occurring in the CHT at 120hpf. *kdrl* (blue), mitophagy (magenta), healthy mitochondria (green) (D): qPCR analysis of *bnip3lb* in *foxo3b*-morphant embryos at 48hpf. Mean ± SEM, n = 4 (E): Image showing mitophagy is increased in the CHT region at 48hpf in prdx1 morphant compared to control. Healthy mitochondria (green), mitophagy (magenta), *kdrl* (blue) (F): Ratiometric analysis of mCherry/GFP in kdrl+ cells in confocal images shows *prdx1* knockdown induced an increase in mitochondria undergoing mitophagy. Mean ± SEM, n = 5 p = 0.0347 (G): qPCR analysis of *bnip3lb* in *prdx1*-morphant embryos at 48hpf. Mean ± SEM, n = 3 (H): Image of ROS in *runx1* and *kdrl* positive HSPCs at 48hpf in the CHT region. *runx1* (magenta), ROS/HyPer (green), *kdrl* (blue) (I): qPCR analysis of *bnip3lb* in *hif1a*-morphant embryos at 48hpf. Mean ± SEM, n = 5

**Supplemental Figure 4:**
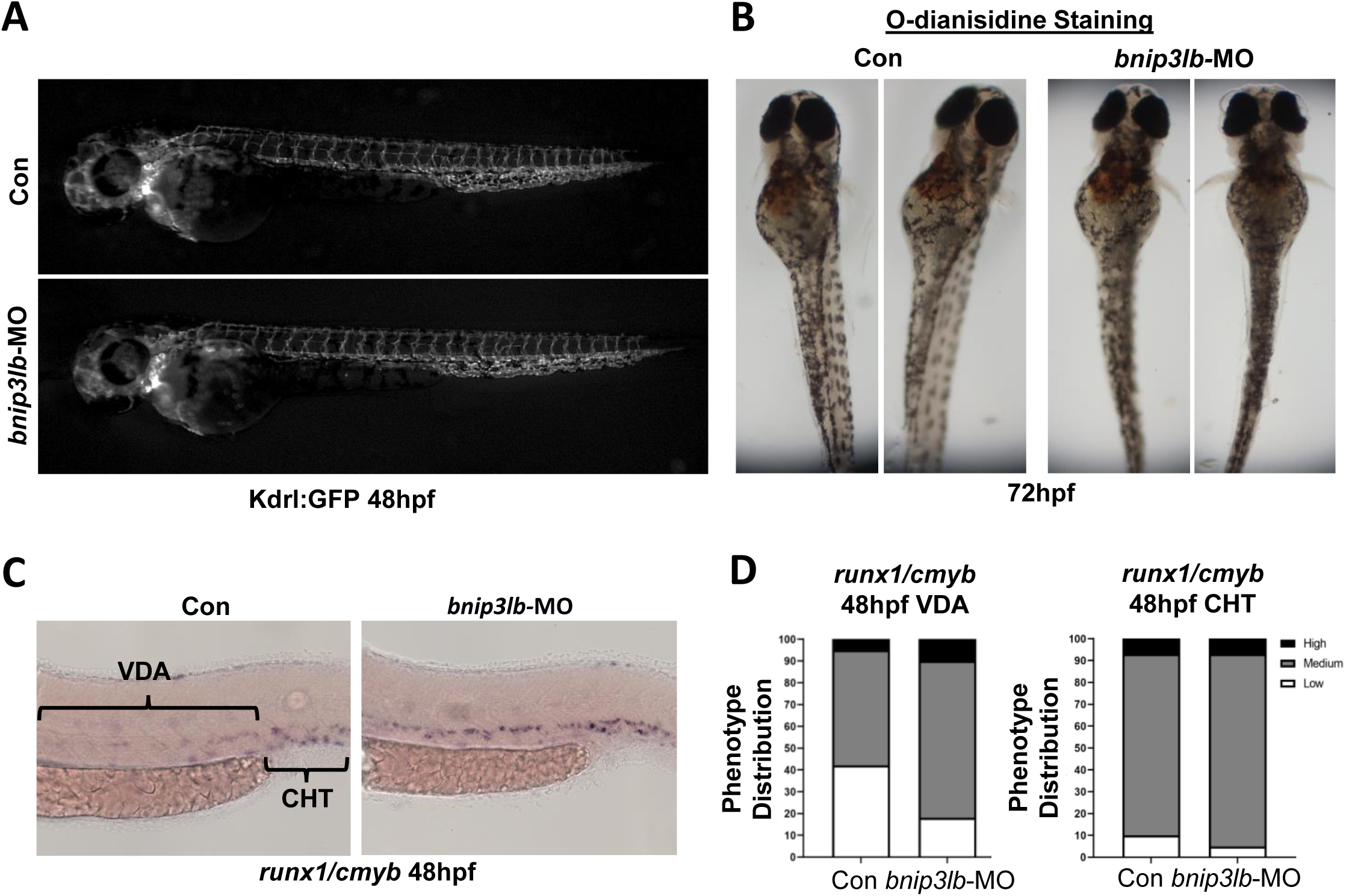
Bnip3lb knockdown does not induce changes in kdrl expression or red blood cell development (Related to Figure 4). (A): Image of kdrl:eGFP vasculature after *bnip3lb* knockdown (B): Image of red blood cells stained with O-dianisidine after *bnip3lb* knockdown (C): Representative images of expression of *runx1/cmyb* after *bnip3lb* knockdown assessed by WISH at 48hpf in the VDA (left) and CHT (right) (D): Phenotypic distribution plot of *runx1/cmyb* expression scored in embryos from (C) (n = 20 per clutch, 3 clutches)

**Supplemental Figure 5:**
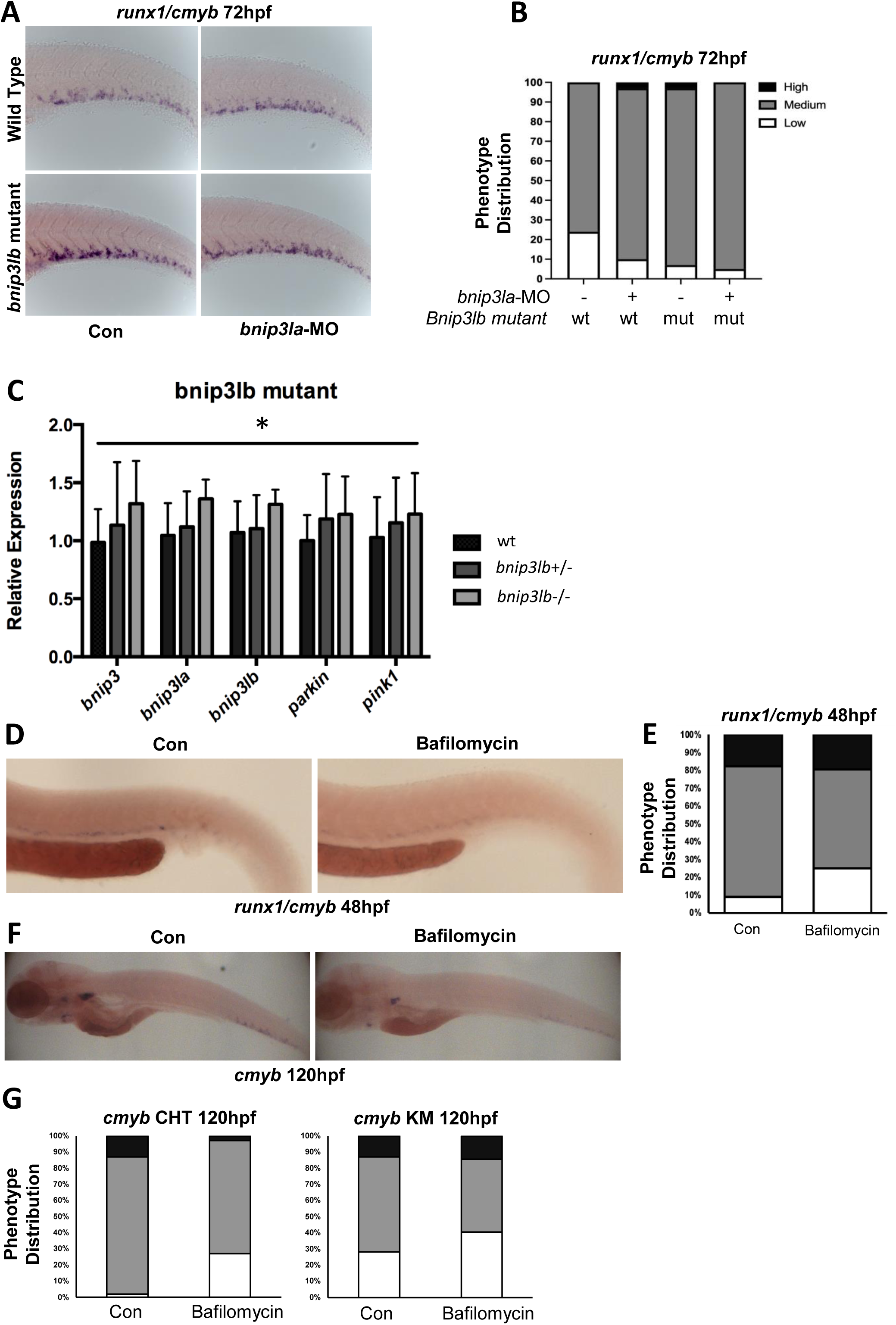
Bnip3lb mutants show signs of genetic compensation but inhibition of autophagy with bafilomycin inhibits HSPC marker expression in the CHT. (A): Representative images of expression of *runx1/cmyb* in control and *bnip3lb* mutant fish after *bnip3la* knockdown assessed by WISH at 72hpf (B): Phenotypic distribution plot of *runx1/cmyb* expression scored in embryos from (A) (n = 20 per clutch, 3 clutches) (C) qPCR data showing wild type vs bnip3lb heterozygous mutants vs bnip3lb homozygous mutants. Mean ± SEM, n = 3, ANOVA used to compare control vs +/- mutants (not significant) and control vs -/- mutants (significant). (D): Representative images of expression of *runx1/cmyb* in control and bafilomycin treated (50mM 24-48hpf) fish assessed by WISH at 48hpf (E): Phenotypic distribution plot of *runx1/cmyb* expression scored in embryos from (D) (n = 20 per clutch, 3 clutches) (F): Representative images of expression of *cmyb* in control and bafilomycin treated (50mM 48-120hpf) fish assessed by WISH at 120hpf (G): Phenotypic distribution plot of *cmyb* expression scored in embryos from (F) (n = 20 per clutch, 3 clutches)

**Supplemental Figure 6:**
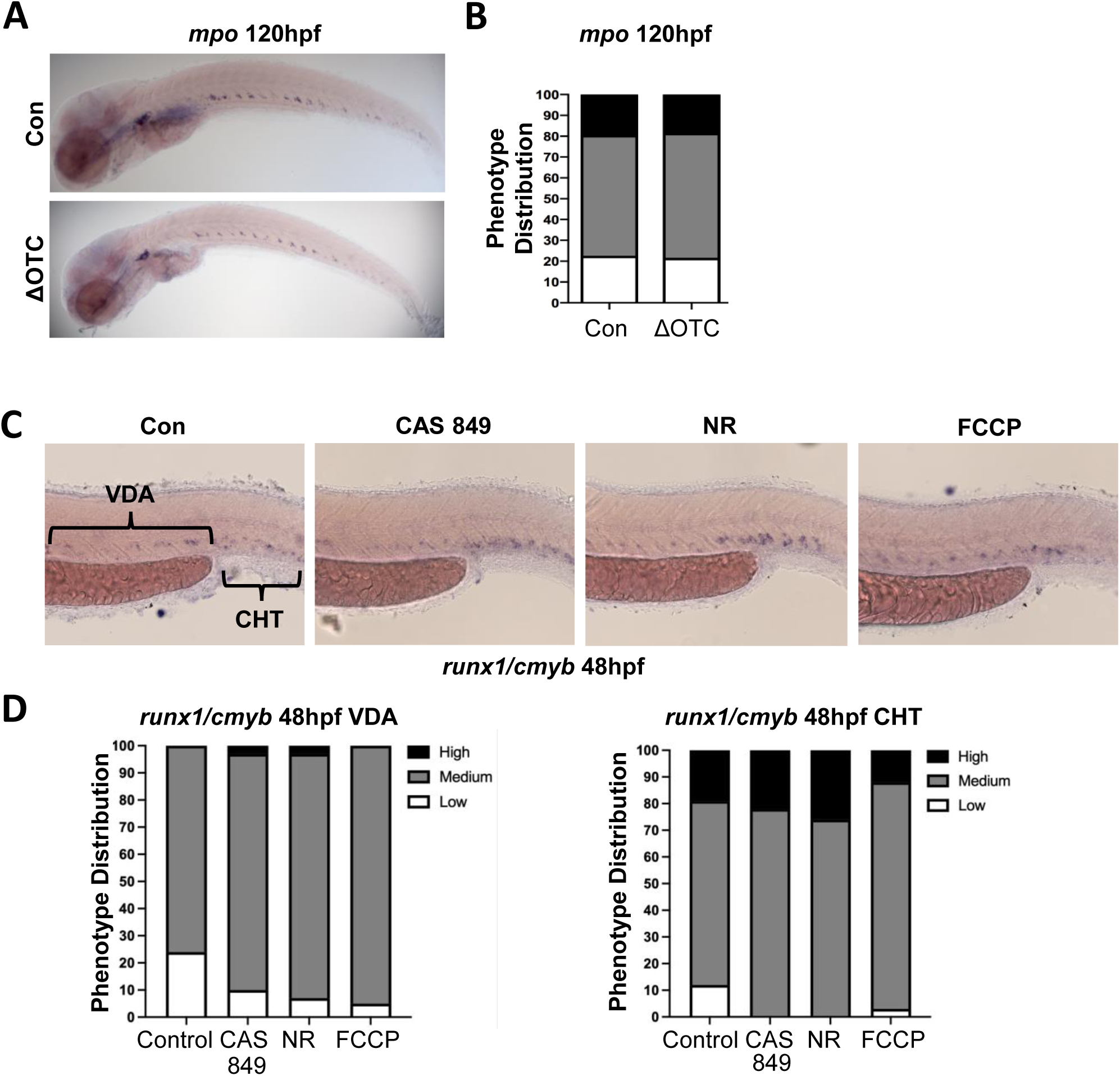
Inducing mitophagy increases HSC numbers significantly in the CHT at 48hpf (Related to Figure 6) (A): Representative WISH images of *mpo* expression in the CHT and kidney marrow at 120hpf, after daily overexpression of *ΔOTC* beginning at 48hpf. (B): Phenotypic distribution plot of *mpo* expression scored in KM of embryos from (H) (n = 20 per clutch, 3 clutches) (C): Images showing expression of *runx1/cmyb* in embryos treated with CAS 849, NR or FCCP (24hpf-48hpf) to pharmacologically increase mitophagy assessed by WISH at 48hpf (D): Phenotypic distribution plot of *runx1/cmyb* expression scored in embryos from (A) (n = 20 per clutch, 3 clutches)

**Supplemental Figure 7:**
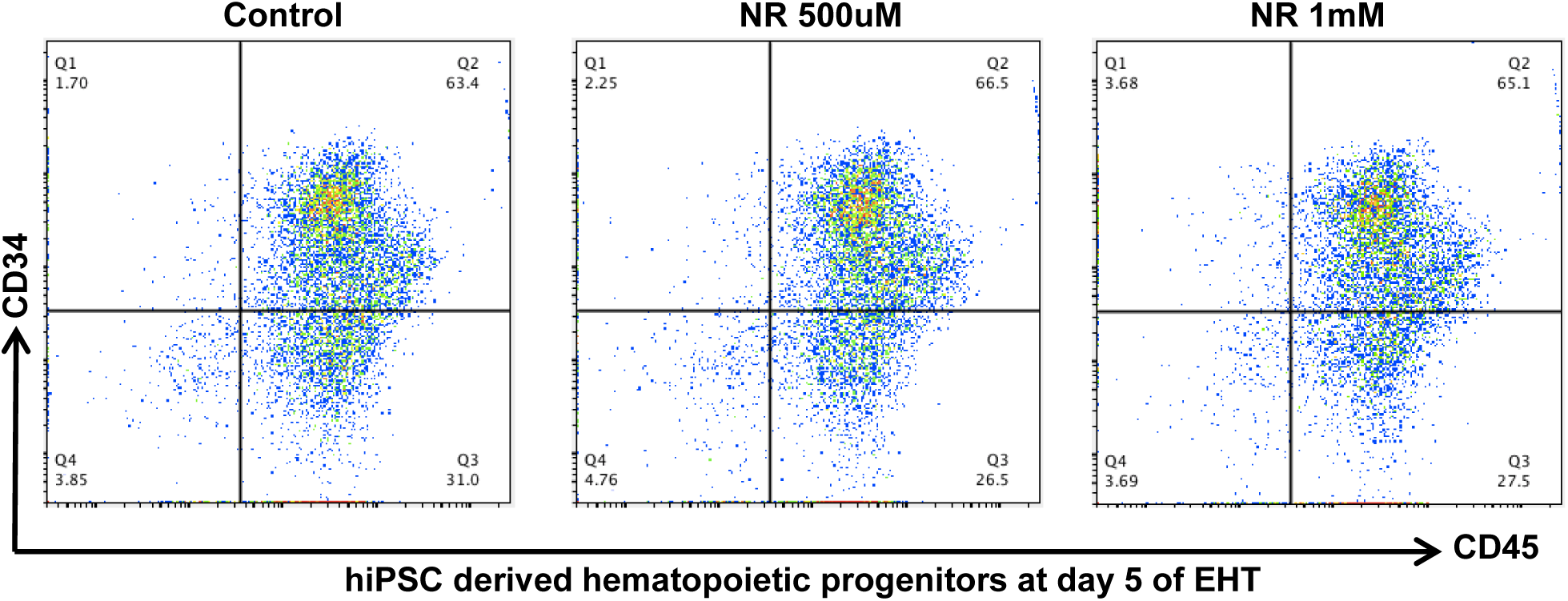
Representative flow plots showing NR treatment results in little change in hiPSC derived CD34+CD45+ HSPCs at day 5 (EHT stage).

**Table S1:**
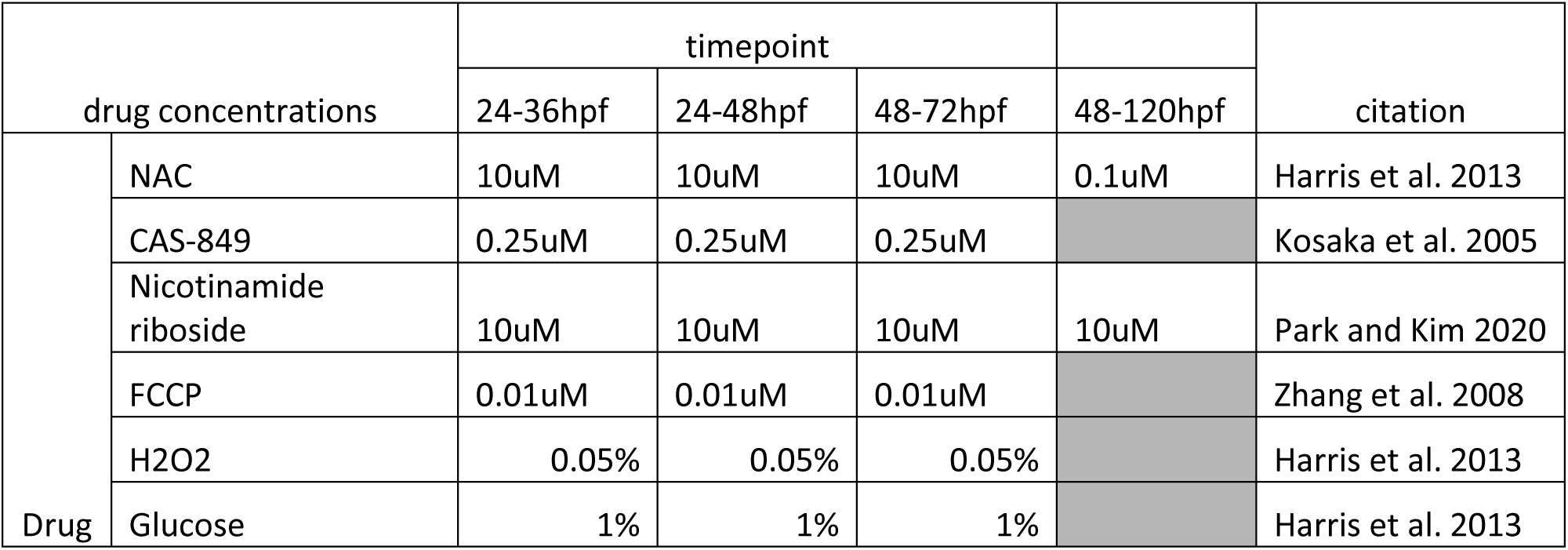

| drug concentrations |  | timepoint |  |  | 48-120hpf | citation |
| --- | --- | --- | --- | --- | --- | --- |
|  |  | 24-36hpf | 24-48hpf | 48-72hpf |  |  |
| Drug | NAC | 10uM | 10uM | 10uM | 0.1uM | Harris et al. 2013 |
|  | CAS-849 | 0.25uM | 0.25uM | 0.25uM |  | Kosaka et al. 2005 |
|  | Nicotinamide riboside | 10uM | 10uM | 10uM | 10uM | Park and Kim 2020 |
|  | FCCP | 0.01uM | 0.01uM | 0.01uM |  | Zhang et al. 2008 |
|  | H2O2 | 0.05% | 0.05% | 0.05% |  | Harris et al. 2013 |
|  | Glucose | 1% | 1% | 1% |  | Harris et al. 2013 |

**Table S2:**
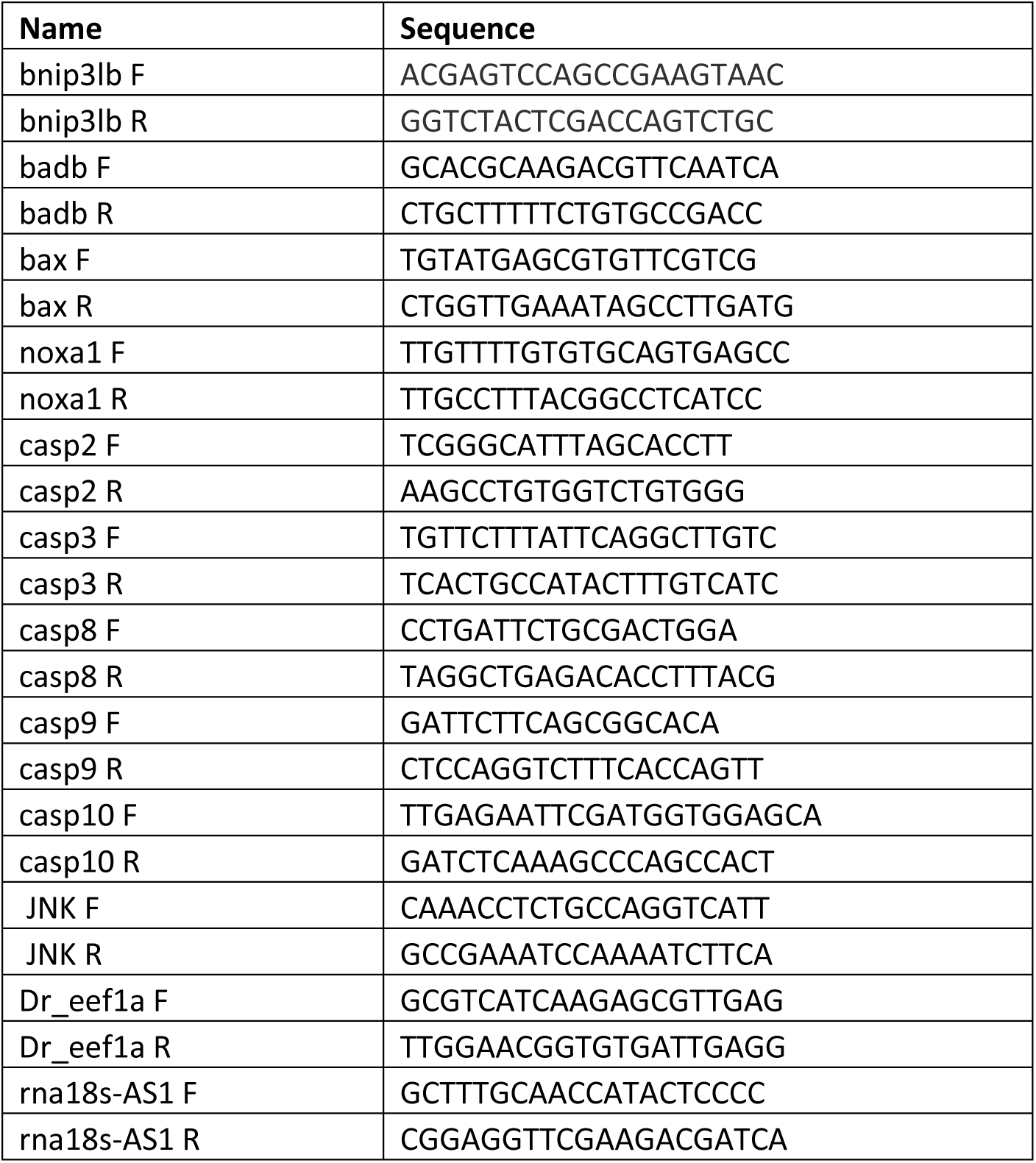

